# The social structure of a captive group of Rüppell’s vultures (Gyps rueppellii)

**DOI:** 10.64898/2026.10.09.757993

**Authors:** Eva S.J. van Dijk, Sophia Wolfswinkel, Jessica H. Smith, Jade A. Zanutto, Aimée Bakker, Zofia Trafas, Maartje H. L. Hooymans, Lisa-Claire Vanhooland, Jorg J.M. Massen

**Author notes:** Corresponding authors: Eva S.J. van Dijk ^a,b,c^,; Jorg J.M. Massen ^a,b,c^. CRediT authorship contribution statement: Conceptualization: E.S.J.v.D., L.C.V., J.J.M.M.; Data curation: E.S.J.v.D., S.W., J.H.S., J.Z., A.M., Z.T., M.H.L.H.; Formal analysis: E.S.J.v.D.; Investigation: E.S.J.v.D.; Methodology: E.S.J.v.D., L.C.V., J.J.M.M.; Supervision: J.J.M.M.; Visualization: E.S.J.v.D.; Writing — original draft: E.S.J.v.D.; Writing — review & editing: E.S.J.v.D., S.W., J.H.S., J.Z., Z.T., M.H.L.H., L.C.V., J.J.M.M.

## Abstract

Studies on the evolution of intelligence have so far mainly focused on mammals, primates in particular, and only few avian taxa, mainly corvids and parrots. Vultures, however, have relatively large brains too, and those of the *Gyps* genus are highly social. Their obligate scavenging lifestyle and sociality make them an excellent taxon for investigating the role of social and ecological selective pressures on the evolution of intelligence. However, little is known of *Gyps* vultures, particularly regarding their social complexity (sensu Social Brain Hypothesis). To investigate the complexity within their social structure, we studied the dominance hierarchy and social bonds of a captive group (n = 23) of Rüppell’s vultures (*Gyps rueppellii*) over two consecutive years. Our results revealed that dominance ranks and affiliative bonds were stable, although there were some differences between breeding and non-breeding seasons. During the breeding season, pair-bonded individuals were more dominant and less social than unpaired individuals. Pair-bond status, age and season influenced dyadic bond strength. Most notably, pair-bonded dyads had substantially stronger bonds, but maintained multiple other strong bonds, too. Our results highlight that multiple forces shape the dominance hierarchy and affiliative network of captive Rüppell’s vultures. Individual characteristics shape social bonds, social relationships are differentiated, and changes between the breeding and non-breeding season can result in social uncertainty, adding to the complexity of this system. The complexity of Rüppell’s vultures’ social structure seems comparable to that of some social corvid and parrot species, where it has been linked to their sophisticated cognition. Future socio-cognitive work on this and other less social vulture species could further identify the social challenges they face, and reveal whether a complex social environment may have been a selection pressure on the evolution of intelligence, both in general, and in this promising new taxonomic group for cognition studies specifically.

## INTRODUCTION

The social intelligence hypothesis (SIH) predicts that cognitive skills emerge from the selective pressures of living in a complex social environment (e.g. Jolly, 1966; Humphrey, 1976; Dunbar and Shultz, 2007), where individuals interact with many others across different contexts and over time (Freeberg et al., 2012). As individuals need to both cooperate (cf. Vygotskian intelligence: Moll and Tomasello, 2007) and compete (cf. Machiavellian intelligence: Byrne and Whiten, 1988) with conspecifics, selection may have favoured those able to solve these social challenges, resulting in the evolution of complex cognitive abilities (Cheney and Seyfarth, 2007). However, according to the ecological intelligence hypothesis (EIH), selective pressures associated with features of the diet have shaped the evolution of intelligence instead (e.g. Parker and Gibson, 1977; Milton, 1981; Sol et al., 2005). Specific cognitive skills may have evolved independently as a result of specific ecological and/or social selective pressures (Allen, 2017; Bräuer et al., 2020), the SIH and EIH vary in explanatory value for cognitive skills in different domains (Rosati, 2017). The SIH was originally formulated to explain primate cognition as primates are highly social, but a recent meta-analysis using 103 studies on 17 different taxonomic orders also finds support for it as explanation for cognitive variation (Speechley et al., 2024). Still, studies testing the SIH predominantly focus on mammals, especially primates and rodents. However, our closest living relatives can only inform us about homologies (Van Horik et al., 2012; cf. Massen, 2021), and without specific knowledge about the common ancestors, not so much about the potential social selective pressures. Thus, to identify those selective pressures that facilitated the evolution of traits such as complex cognition, research should focus on more distantly related taxa where these traits likely evolved independently through convergence. As the amniote lineage (includes birds, reptiles and mammals) split approximately 310-320 million years ago (Damas et al., 2022; Deakin and Ezaz, 2014), it is likely that any observed more complex cognitive skill of birds and mammals arose independently, making birds an interesting group for studying the evolution of cognition and testing the SIH.

Research on the link between social complexity and cognition in birds has so far, with a few notable exceptions (e.g. Weiß et al., 2010; Ashton et al., 2018), mainly focussed on corvids and parrots. Both parrots and corvids are long-lived, highly social birds with long-term monogamous pair-bonds and altricial young that are dependent for extended periods (Emery, 2005; Lambert et al., 2019). They are also suggested to have complex social structures (e.g. Bugnyar, 2024), for example individuals form multiple individualized relationships and use social information in decision-making processes (Boucherie et al., 2019; Penndorf et al., 2025, 2026b). Moreover, they stand out from other bird species as their intellectual abilities are consistently more elaborate, and in many domains comparable to those of monkeys and apes (e.g. Lambert et al., 2019; Pika et al., 2020). Following the SIH, the difference in cognitive abilities between parrots and corvids and other birds could be the result of their sociality (Boucherie et al., 2019; Emery, 2005), but it could also be due to the limited number of bird species tested. Pigeons (*Columba livia*), for example, perform on par with corvids and primates on some tasks (Güntürkün et al., 2017). Thus, to better understand the evolution of avian cognition, and to determine the generalizability of the SIH (Speechley et al., 2024), it is vital that the cognitive abilities of a wider range of taxonomic groups are studied in relation to their socio-ecology.

One interesting taxonomic group for the study of avian cognition in relation to socio-ecological factors is vultures. While parrots and corvids are genetically distant from vultures (Jarvis et al., 2014; Prum et al., 2015; Stiller et al., 2024; Wink et al., 2026), there are similarities when it comes to their socio-ecology. Vultures of the *Gyps* genus, in particular, resemble parrots and corvids, as they are also highly social, form long-term monogamous pair bonds, and have a slow life history (Emery, 2005; Lambert et al., 2019; van Overveld et al., 2022). However, where parrots and corvids are omnivorous generalists, *Gyps* are highly specialized obligate carrion consumers that rely on aerial social networks for finding food (Bildstein, 2022; Cortés-Avizanda et al., 2014; van Overveld et al., 2022). The similarity of *Gyps* sociality to that of parrots and corvids, combined with the challenge posed by their feeding ecology (cf. EIH: Parker and Gibson, 1977; Milton, 1981; Sol et al., 2005; Rosati 2017), could allude to the evolution of elaborate cognitive skills in social vultures.

While there has been little in-depth and/or experimental research on *Gyps* vultures’ cognitive abilities, the limited evidence so far suggests that they do indeed possess relatively elaborate cognitive skills and have relatively large brains (van Overveld et al., 2022). Griffon vultures (*Gyps fulvus*), the most studied *Gyps* species, rely on both environmental and social information to navigate to the next updraught (Sassi et al., 2024). Informed griffon vultures are also followed by uninformed individuals from the roost to food sources (Acácio et al., 2024; Harel et al., 2017). They may use such social information flexibly and strategically, as lower ranking individuals are more likely to disregard social cues, potentially to avoid their competitors (Bosè et al., 2012; Sassi et al., 2024). In addition, it has been suggested that they show complex fission-fusion dynamics while soaring (van Overveld et al., 2022). Griffon and Rüppell’s vultures (*G. rueppellii*) also have an articulated vocal repertoire, which would suggest that they have a complex social system that requires such a complex communicative system (cf. “The Social Complexity Hypothesis for Communicative Complexity”: Freeberg et al., 2012; Peckre et al., 2019), as well as the cognitive skills to maintain it (Romani et al., 2022; Proudfoot et al., in press). As the SIH predicts that a complex social environment is a key selective pressure shaping the evolution of cognitive skills, and *Gyps* vultures may possess relatively elaborate socio-cognitive abilities, these vultures may have a complex social system.

To gain a better understanding of the evolution of the cognitive abilities of vultures and to test the SIH, the social system of *Gyps* vultures must first be mapped so that the potential complexity within this system can be identified (Kappeler, 2019). According to Kappeler (2019) a social system is made up of four main components; i. The social organization component refers to group size and composition, ii. the mating system to mating and reproduction patterns and iii. the care system to parental care and iv. the social structure to the content, quality, and patterning of social relationships, which in turn result from the interactions among individuals. These components of a social system directly influence the nature and patterning of social interactions, and thus the social challenges individuals face. As such, both the overarching social system and an individual-level perspective are important when investigating social complexity (Boucherie et al., 2019).

When applying the social system framework (Kappeler, 2019) to what is known of *Gyps* vultures, it becomes apparent that the mating and care system and the social organisation of most species have been documented in the wild (e.g. Mundy et al., 1992; Xirouchakis, 2007; Xirouchakis and Mylonas, 2007; Yadav Ruby and Kanaujia, 2018). However, little information is available on their social structure, as this is much harder to grasp without structured and long-term behavioural studies on known individuals. Two aspects that can be used to quantify social complexity, relationship differentiation and social uncertainty, specifically relate to social relationships (Hobson et al., 2019; Kappeler et al., 2019). Social systems become increasingly complex with a greater diversity in social relationships, referred to as relationship differentiation, and when the social environment and interactions become less predictable, also known as social uncertainty. Thus, uncovering the social structure of these vultures is highly relevant when investigating the potential complexity of their social environment.

Social relationships such as dominance relationships and affiliative bonds make up the social structure component of social systems (Kappeler, 2019). Dominance relationships arise from repeated agonistic interactions, such as the conflicts that *Gyps* experience around carcasses (Kane et al., 2022; Sassi et al., 2024). That these conflicts rarely escalate to injury (Mundy et al., 1992) could indicate that there is a dominance relationship among individuals that regulates the access to resources and reduces aggression (Tibbetts et al., 2022). Relationships that instead have relatively low rates of agonism and high rates of affiliative interactions are called (affiliative) social bonds (Kappeler, 2019; Massen et al., 2010). However, even within their long-term breeding pairs, they display little physical contact and they are therefore considered undemonstrative partners (Mundy et al., 1992; Xirouchakis and Mylonas, 2007). This absence of overt affiliative behaviour makes it difficult to identify social bonds when observing colonies of wild *Gyps.* Nevertheless, potential evidence of differentiated affiliative bonds among wild *Gyps* comes from griffon vultures focussing on spatial proximity patterns (cf. Massen et al., 2010), which shows that the distance between informed individuals and followers in flight on foraging trips varies structurally, suggesting a tendency to fly with or to avoid specific conspecifics and therefore variation in social bond strength (Acácio et al., 2024; Harel et al., 2017). These patterns also do not necessarily restrict themselves to pair-bonded partners, as for example in a captive colony of Cape vultures (*G. coprotheres*), paired individuals are reported to sometimes form a relationship with an “extra-pair partner” during the breeding season (Hirschauer and Wolter, 2017). Thus, the social bonds of *Gyps* vultures may not be limited to the pair-bond but could also include dominance relationships, extra-pair partnerships and other social bonds.

To fill in some of the gaps in our understanding, we aimed to do an in-depth investigation of the social structure of a captive population of Rüppell’s vultures. In captivity, group size and composition are the result of husbandry decisions, but as individuals can still form and maintain social relationships, relationship differentiation and social uncertainty could still be informative measures of social complexity as experienced by individuals (Hobson et al., 2019; Kappeler et al., 2019).

Specifically, we investigated dominance ranks and social bonds of a captive group of 23 Rüppell’s vultures of varying ages and sexes, including five pairs (i.e., 10 pair-bonded individuals), for two consecutive years. We tested whether *dominance ranks* (Research question (RQ) 1) and hierarchy steepness (RQ 2) were stable over time and whether individual characteristics or season predicted individual dominance ranks (RQ 3). For the *social bonds*, we investigated whether individual sociality (RQ 4) and the number of strong bonds (RQ 5) was related to individual characteristics and season. On a dyadic level, we also tested whether social bonds were stable over time (RQ 6) and whether they were related to individual characteristics and differences in dominance rank (RQ 7).

In a small captive mixed group of griffon and Rüppell’s vultures the dominance hierarchy was steep and consistent between two years (Sassi et al., 2024), thus we also expected a steep and stable dominance hierarchy in the captive population we studied (RQ 1 & 2). Neither sex nor age predicted rank in this group (Sassi et al., 2024), but in wild griffon vultures, dominance ranks were considered higher in old adults (Bosè and Sarrazin, 2007). We thus expected dominance rank to be predicted by age, with older individuals outranking younger individuals (RQ 3). In wild griffon vultures, age-related effects on social networks have also been found, as older individuals tended to be more selective in their social bonds, especially during the breeding season (Acácio et al., 2024). We thus expected older individuals, and especially pair-bonded individuals during the breeding season, to be less social (RQ 4) and have fewer strong bonds (RQ 5 & 7). Despite seasonal fluctuations in individual sociality (Acácio et al., 2024), we expected dyadic affiliative bonds to be relatively stable over the two years (RQ 6). As younger individuals may be more social, we expected their dyadic bonds to be stronger (RQ 7). In non-breeding aggregations of ravens (*Corvus corax*), individuals are suggested to prefer bonding with higher-ranking individuals (Braun and Bugnyar, 2012), thus, we also expected that increased rank differences may increase bond strength. Finally, as Rüppell’s vultures form lifelong pair-bonds, we expected pair-bonds to be the strongest affiliative bond (RQ 7).

## METHODS

### Subjects and Housing

We studied a group of Rüppell’s vultures housed in Avifauna bird park, Alphen aan de Rijn, the Netherlands (n = 23, 9F, 5 pairs) (Supporting Online Information (SOM) table S1). Individuals were recognizable using coloured leg bands. As exact ages of ten individual were unknown, we assigned individuals into age classes similar to those used in studies of wild griffon vultures based on their age at the start of each observation period (SOM table S1) (Acácio et al., 2024; Bosè and Sarrazin, 2007). From hatching until the end of their first breeding season, individuals are referred to as chicks. Individuals were only included as subject from their first non-breeding season onwards, i.e. after they fledged, were about half a year old, and started interacting with the rest of the group. These young individuals were referred to as immatures from this first non-breeding season onwards until 4 years old (YO). Young adults are between 5-10 YO and adults are over 10 YO. The group size ranged from 16 to 23 individuals as seven chicks hatched during the study, while two immatures were moved out of the group. To increase hatching success, eggs are replaced with dummy eggs and incubated artificially. Newly hatched chicks are returned a few days after hatching to a nest with either biological or foster parents.

The vultures were housed in a ±14×34×4.5m aviary with nest boxes, a variety of natural perches and ad libitum access to fresh water. They were fed a variety of carrion items, including large pieces of cow carcass and whole rabbits. Feedings were typically in the late afternoon every one to three days depending on the nutritional needs of the birds, similar to feeding patterns in wild *Gyps* populations (e.g. Bosè and Sarrazin, 2007). The aviary also housed a pair of pied ravens, and during some brief periods two white-headed vultures, one or two hooded vultures, and/or one or two Southern ground hornbills.

### Ethical Note

This study was conducted at Avifauna Bird Park, an accredited member of the European Association of Zoos and Aquaria (EAZA) where the vultures are cared for in accordance to EAZA’s Standards for the Accommodation and Care of Animals in Zoos and Aquariums (EAZA 2024). The data was collected noninvasively through observations that were done from outside of the enclosure.

### Data Collection

Data were collected from March through June and September through December 2023 and for all of 2024, thus covering 20 months, including both breeding (January-June) and non-breeding (July-December) seasons. For clarity, the two breeding and two non-breeding seasons during which data was collected are referred to as four (observation) periods.

To determine the dominance hierarchy, we recorded a feeding (“food monopolization observations”) about every two weeks (cf. Boucherie et al., 2022). During these observations, food was provided in a single location by a caretaker and the first 30 minutes of interactions around the food were filmed in a wide angle to capture as many agonistic interactions as possible. Both contact aggression and non-contact agonistic behaviours were recorded using ad libitum all-occurrence sampling in B.O.R.I.S. (Friard and Gamba, 2016), noting the aggressor, recipient and outcome of each interaction where individual identities could be determined (SOM table S2).

Coding was split between three different coders, E.S.J.v.D., S.W., and A.B., with a good inter-rater reliability (ICC(A,1) = 0.784, 95%CI [0.746;0.817]) (Koo and Li, 2016) (SOM Appendix S1). A total of 40 feedings were coded, 12 during the breeding and 6 during the non-breeding season of ’23, and 13 and 9 feedings for the respective seasons in ’24. Only agonistic interactions with a clear winner (i.e., unidirectional) were used to determine the dominance hierarchy, referring to interactions where the recipient responded with a submissive signal, retreated or was chased away.

To estimate the social bonds of Rüppell’s vultures, we collected 10-minute focal follow observations. In total, 3233 focal observations (572 hours, mean = 7 hours 9 minutes per subject per period, range 2 hours 3 min. – 16 hours 24 min.) were collected, 478 during the breeding and 601 during the non-breeding season of ’23, and 1438 and 716 during the respective seasons in ’24 (SOM Table S4). As other affiliative behaviours such as allo-preening and contact-sitting were rare, and spatial proximity has previously been used to estimate social bonds in captive *Gyps* vultures (Sassi et al., 2024), we here focused on proximity.

Coding of focal observations was conducted by E.S.J.v.D., S.W., J.H.S., J.Z., Z.T., A.B. and M.H.L.H. using B.O.R.I.S., scoring social proximity (within one wing-length) and, for pair-bonded individuals, the time they spent on their nest alone or together (SOM Table S3-S4) (Friard and Gamba, 2016). An interrater reliability assessment including all coders apart from J.H.S. and J.Z. showed good agreement (ICC(A,1) = 0.792, 95%CI [0.73, 0.847]) (Koo and Li, 2016) (SOM Appendix S1.). J.H.S. and J.Z. could not be compared directly as they did not code the same observations, so to assess their reliability, they were each compared to the five other coders. J.Z. achieved good interrater reliability (ICC(A,1) = 0.821, 95%CI [0.741, 0.888]) (Koo and Li, 2016) and J.H.S. reached a moderate interrater reliability (ICC(A,1) = 0.707, 95%CI [0.603, 0.803]).

### Analyses

All analyses were performed using RStudio (Posit team, 2023; R Core Team, 2025). For ease of reference, we numbered all final models. For these models, we checked model diagnostics to see if model assumptions were met (R package DHARMa (Hartig et al., 2024)) (SOM Appendix S2. SOM Figures S1-S10). In addition, when model outputs included significant effects of predictors with more than one level and/or significant interaction effects, we performed Tukey post-hoc pairwise comparisons (R package emmeans (Lenth et al., 2026)).

### Analyses Dominance Hierarchy

#### Dominance rank determination

To investigate the predictors of dominance rank and the stability of hierarchy steepness, agonistic interactions with a clear winner were used to calculate Bayesian Elo ratings (R package EloSteepness (Neumann, 2023)) (Neumann and Fischer, 2023). This was done for each period separately, only including individuals present during the whole period (n = 16-21). On average 115 agonistic interactions per subject per period were recorded, surpassing the recommended minimum of 10 interactions (Sánchez-Tójar et al., 2018). To calculate the Bayesian Elo ratings, we ran 4 chains of 2000 iterations with a warm-up of 1000 iterations for each of the four periods. This gave us posterior distributions of the individual Elo-ratings, which are individual summed winning probabilities (SOM Table S5.).

Individual ordinal ranks were determined based on the mean Elo-ratings, where the individual with the highest Elo-score within a period was assigned the highest rank. Finally, for each period, we also inferred the hierarchy steepness, which is the slope of the regression of the summed winning probabilities as a function of the ordinal ranks (Neumann and Fischer, 2023). This value can range from 0 to 1 and reflects the absolute difference in the overall winning success of adjacently ranked individuals (de Vries et al., 2006).

#### Dominance rank stability

To determine the stability of the dominance hierarchy, all interactions were first used to infer daily Elo-ratings (R package EloRating (Neumann and Kulik, 2024)). Elo ratings reflect the ability of an individual to win a subsequent interaction and thus individual dominance. All individuals present at the beginning of data collection started with an Elo score of 1000 and new individuals (n = 7) were assigned the lowest score on the day they entered (see methods Smit and Robbins, 2025). The two individuals that were moved out of the group, were only included until this date. Next, the S index was calculated (Neumann and Kulik, 2024), which ranges between 0 and 1 and reflects the overall hierarchy stability (RQ 1) (McDonald and Shizuka, 2013; Neumann et al., 2011). A hierarchy is considered stable when individuals maintain the same relative rank position over time. An S index of 1 reflects the absence of rank changes and thus a perfectly stable hierarchy.

#### Stability hierarchy steepness

While we investigated the stability of the dominance rank by calculating the S index based on rank reversals, we were also interested in whether the steepness of the hierarchy would differ between the breeding and non-breeding season. To test this, we compared the steepness of the two breeding and two non-breeding seasons using a Welch t-test (RQ 2).

#### Predictors of dominance

To test whether individual characteristics predict individual dominance rank, we tested the effects of age class, sex, season, having a pair-bond and the interaction between season and pair-bond status on Elo-scores using a generalized linear mixed model (GLMM) with a Gaussian log link family (R package glmmTMB (Brooks et al., 2026)). We included random effects for individual identity and period to account for repeated measures and temporal variation (RQ 3). While each pair (n = 5) incubated an (artificial) egg in both breeding seasons and raised a chick in 2023, only three pairs raised a chick in 2024. As there could be behavioural differences between pairs with and without dependent offspring in the breeding season, we first tested whether a model where we differentiated between pairs with and without a chick during the breeding season would have a significantly better fit than a model only including paired or non-paired as levels. These models were compared using ANOVA. Separating pairs with and without dependent offspring did not improve model fit (ANOVA: χ^2^ =0.139, *P =* 0.709), thus the simpler model with only two levels for pair-bond status was selected for further analyses (**model 1**).

### Analyses Social Bonds

#### Social bond estimation

We used the time a dyad spent in proximity as a proxy for their social bond strength (cf. Massen et al., 2010). First, to prevent underestimating the bonds between pair-bonded and un-paired individuals, we omitted the observational data collected while the focal individual was on the nest, as it is unlikely that a non-paired individual would be allowed in proximity of its nest (griffon vultures: Xirouchakis and Mylonas, 2007) (SOM Table S3-S4). For each dyad, we determined the time they spent in proximity and the total time the two individuals were observed. With these measures, we calculated a symmetric proximity rate (minutes per hour observed). We then generated undirected, weighted social networks for each period in order to calculate individual strength, which reflects the total sum of proximity rates (m/h) per sociality (Brent et al., 2011). To investigate the extent of relationship differentiation, we focused on the number of strong bonds per individual. Dyads that spend more time together in proximity than the group average for that period were considered to have a strong bond (McFarland et al., 2017; Schülke et al., 2022). We then counted the number of strong bonds per individual per period. Finally, we used the dyadic proximity rates as a proxy for dyadic social bond strength to investigate what shapes social bonds. If a dyad was not observed in proximity, they were included as actual zeros in the social bond strength analyses, as their true time in proximity was likely very low based on the amount of focal data collected.

#### Predictors of sociality

Next, we investigated whether individual characteristics and season influence individual sociality as measured through strength (total proximity rate (m/h)) and eigenvector centrality (Brent et al., 2011). In our model we included age class, sex, z-transformed Elo score and an with minor deviations in the upper quantile for eigenvector centrality (SOM Figure S3-S6).

#### Relationship differentiation

To investigate whether social bonds were differentiated and whether pair-bonded individuals maintained strong affiliative bonds outside of the pair-bond, we studied what predicted the number of strong bonds individuals maintained. To this end, we modelled the number of strong ties using a GLMM with a gamma log link, and included age class, sex and an interaction between pair-bond status and season as predictor variables, and period and individual identity as random effects (**model 4**). To meet model assumptions, we dropped individuals for those periods where we could identify no strong ties, which corresponds to them having spent less time in proximity with their interaction partners than the group average (McFarland et al., 2017; Schülke et al., 2022). Based on this criterium, two individuals were dropped from a single breeding season during which they had a dependent chick. Model diagnostics suggested a good fit (SOM Figure S7-S8).

#### Stability of social bonds

To determine whether dyadic social bonds remained stable over time, we used a multiple regression quadratic assignment procedure (MRQAP; Krackhardt, 1988; Dekker et al, 2007; see methods Borgeaud et al., 2017). We conducted MRQAPs between matrices of dyadic proximity rates of each period and all periods thereafter. For each MRQAP, we only included individuals that were present during both periods (cf. Massen and Sterck, 2013). We ran 5000 permutations per MRQAP and corrected for multiple testing using the Bonferroni method.

#### Predictors of social bonds

To investigate what shapes the social bonds of Rüppell’s vultures, we investigated the potential predictors of dyadic proximity rates (m/h) (RQ 6). As the dyadic proximity per period consisted of continuous positive values but also contained zeros for dyads that were not observed together, we used a GLMM with a Tweedie distribution. We formulated a model that included age combination, sex combination, an interaction between pair bond and season, and, as a measure of rank difference, the z-transformed absolute difference in Elo scores (**model 5**). We included random effects for dyad identity and period. Model diagnostics suggested a generally good fit (SOM Figure S9-S10).

## RESULTS

### Dominance

To estimate whether a group of Rüppell’s vultures had a stable dominance hierarchy over the course of our study, we calculated an S index from the daily Elo-ratings of 23 individuals. These ratings were inferred from 9318 decided agonistic interactions during the 40 food monopolization trials (mean = 810.3 interactions per trial, median = 654, range: 40 - 1834) observed between 22/03/2023 and 13/12/2024. The S-index was 0.9831, where an index of 1 reflects absolute stability. The dominance hierarchy of Rüppell’s vultures was thus highly stable and rank reversals during that period were rare.

To gain a more detailed understanding of the dynamics underlying the dominance hierarchy structure, we also inferred individual Elo-ratings for each of the four periods separately using a Bayesian approach. We estimated the steepness of each hierarchy based on the individual Elo-ratings (mean steepness = 0.60, range: 0.50 – 0.74) (SOM Table S5) (Neumann and Fischer, 2023). To test if the hierarchy steepness was influenced by season, we compared the steepness of the hierarchy during the two breeding and two non-breeding seasons. The steepness of the Rüppell’s vulture hierarchy was moderately high (e.g. Balasubramaniam et al., 2012) and similar during the breeding and non-breeding season (Welch t-test: t = 1.28, df = 1.52, *P =* 0.363).

To investigate if age, sex, having a pair-bond and/or the breeding season influenced the dominance rank of Rüppell’s vultures, we fitted a GLMM on the Elo scores. Model diagnostics suggested a good fit (SOM Figure S1-S2). Females were more dominant than males, as the Elo scores of males were 28% lower (GLMM: Est. = -.33 ± 0.09, *P <* 0.001, Figure 1A). The effect of having a pair-bond differed between the seasons as we found a significant interaction effect (Est. = 0.40 ± 0.10, *P <* 0.001), warranting post-hoc analyses comparing the effect of pair-bond within each season and the effect of season per pair-bond status using Tukey adjustment for multiple pairwise comparisons. Having a pair-bond influenced dominance during the breeding season, when unpaired individuals had 45% lower Elo scores than pair-bonded individuals (Tukey: Est. = -0.60 ± 0.13, adj. *P <* 0.001, Figure 1B). Moreover, there was an effect of season, as unpaired individuals had 58% higher Elo scores during the non-breeding season than during the breeding season (Est. = 0.46 ± 0.15, adj. *P =* 0.002, Figure 1B). As individual dominance ranks were highly stable, these seasonal effects reflect changes in the competitive ability, so the likelihood of winning an agonistic interaction, of individuals, rather than changes in ordinal ranks. The dominance rank of individuals was thus influenced by sex, pair-bond and season but not by age, with females having higher Elo scores than males, and unpaired individuals having lower scores than pair-bonded individuals during the breeding season (SOM Table S6).

**Figure 1.**
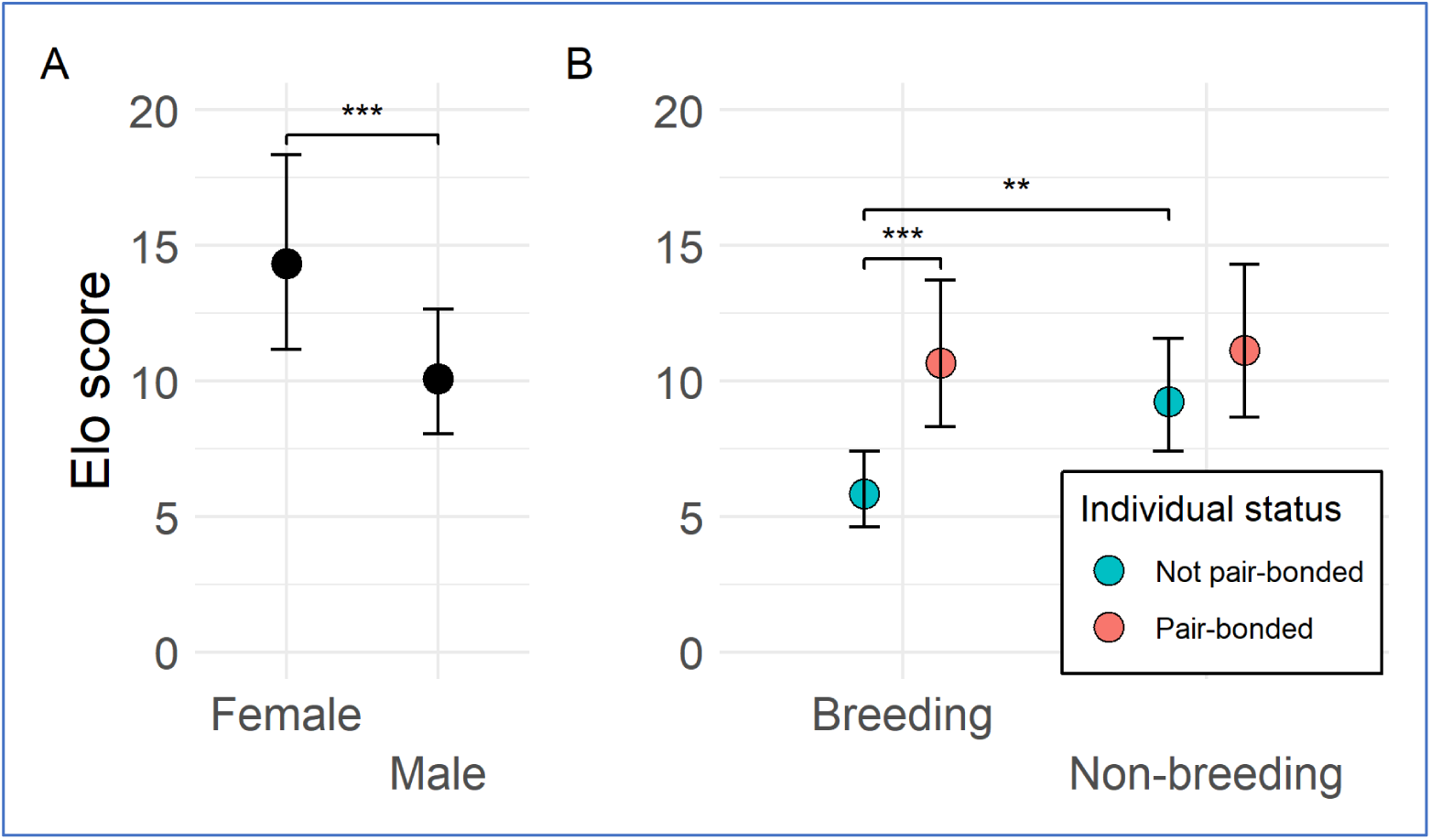
**A)** Marginal effect plot for the predicted Elo scores per sex, averaged over age, season and pair-bond status. Error bars reflect standard error. *** P < 0.001. **B)** Marginal effect plot for the predicted Elo-scores for paired and unpaired individuals during the breeding and non-breeding season, averaged over age and sex. Error bars reflect standard error. ** P < 0.01 and *** P < 0.001 based on Tukey adjusted post-hoc analyses.

### Affiliative Social Bonds

To investigate whether individual characteristics and season influenced individual sociality, we modelled the total proximity rate (m/h) (**model 2**) and the eigenvector centrality (**model 3**) per period. The conditional models both indicated that age had an effect as immatures spent longer in proximity than the adult baseline (**model 2**) (GLMM: Est. = 0.25 ± 0.12, *P =* 0.035) and had a higher eigenvector centrality (**model 3**) (GLMM: Est. = 0.27 ± 0.13, *P =* 0.041). and season only returned a trend indicating that adults may have spent less time in proximity than immatures (Est. = -0.25 ± 0.12, Tukey adj. *P =* 0.089) and no difference in Eigenvector centrality between age classes (Tukey adj. p > 0.05).

Pair-bond status interacted with season to influence both total time in proximity (GLMM: Breeding season : Pair-bonded: Est. = -0.48 ± 0.14, *P <* 0.001, **model 2**) and centrality in the social network (GLMM: Breeding season : Pair-bonded: Est. = -0.46 ± 0.15, *P =* 0.002, **model 3**) (**Figure 2**). During the breeding season, unpaired individuals spent more time in proximity with others than individuals with a pair-bond (Tukey: Est. = 0.35 ± 0.14, adj. *P =* 0.010) and were more central, too (Tukey: Est. = 0.37 ± 0.15, adj. *P =* 0.01). The season also influenced the sociality of pair-bonded individuals, as they spent more time in proximity (Tukey: Est. = 0.31 ± 0.12, adj. *P =* 0.011) and were also more central during the non-breeding season than during the breeding season (Tukey: Est. = 0.25 ± 0.10, adj. *P =* 0.012). For unpaired individuals, the seasons had the opposite effect as they were more central in the breeding season than during the non-breeding season (Tukey: Est. = -0.21 ± 0.10, adj. *P =* 0.038). Thus, pair-bonded individuals were less social during the breeding season than during the non-breeding season, whereas dominance rank and age did not have a significant effect on sociality (SOM Table S7-S8).

**Figure 2.**
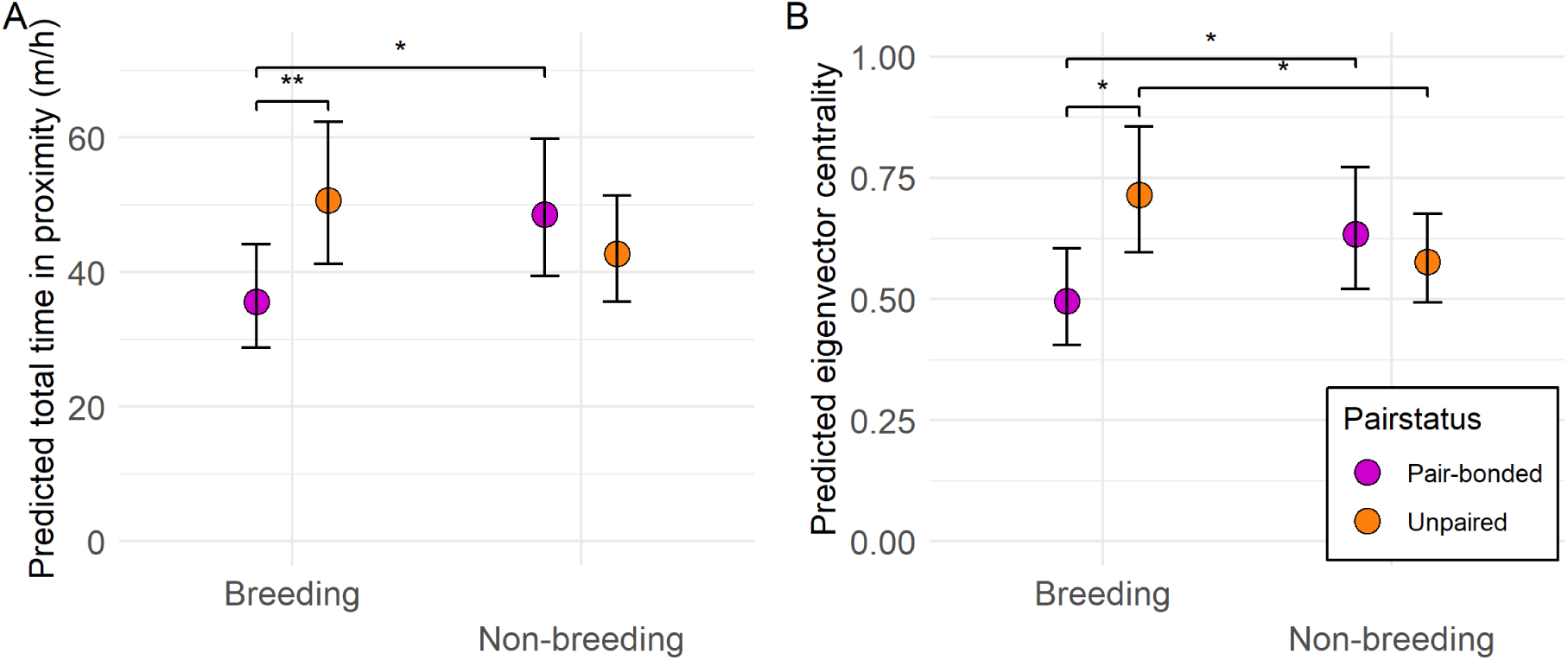
**A)** Marginal effect plot for the predicted total time in proximity off of the nest per individual (strength, m/h) per pair-bond status and season, averaged over age and sex. Error bars reflect standard error. **B)** Marginal effect plot for the predicted eigenvector centrality per pair-bond status and season, averaged over age and sex. Error bars reflect standard error. (* P < 0.05 and ** P < 0.01, Tukey adjusted)

To study relationship differentiation, we investigated whether individual characteristics and season predicted the number of strong bonds. The conditional model suggested an effect for age as both immatures and young adults had more strong ties than the adult baseline (**model 4**) (GLMM: immature Est. = 0.38 ± 0.16, *P =* 0.021; young-adult est. = 0.41 ± 0.19, *P =* 0.032) (**Figure 3**). However, a post-hoc pairwise comparison averaged over pair bond status, sex and season only returned trends suggesting that immatures and young-adults may have had more strong bonds than adults (Tukey: Est. = -0.38 ± 0.16, adj. *P =* 0.055, est. = -0.41 ± 0.19, adj. *P =* 0.082, respectively). A further trend suggested an effect for the interaction between the season and whether an individual had a pair-bond on the number of strong bonds (GLMM: Breeding season: Pair-bonded: Est. = -0.36 ± 0.19, *P =* 0.061, **model 4**). Thus, we performed a post-hoc analysis to investigate the potential effect of season and pair-bond status on strong bonds in more detail, averaged over age and sex. During the breeding season, unpaired individuals had 50% more strong ties than pair-bonded individuals (Tukey: Est. = 0.40 ± 0.17, adj. *P =* 0.017). Nonetheless, pair-bonded individuals maintained strong affiliative bonds outside of their pair-bond as they had more than a single strong bond during the breeding season when estimated over sex and age (95%CI = [4.53;7.48]). During the breeding season, pair-bonded individuals had fewer strong bonds than unpaired individuals whereas sex had no effect on the number of strong bonds (SOM Table S9), with individuals maintaining an average of 6.6 strong ties overall.

**Figure 3.**
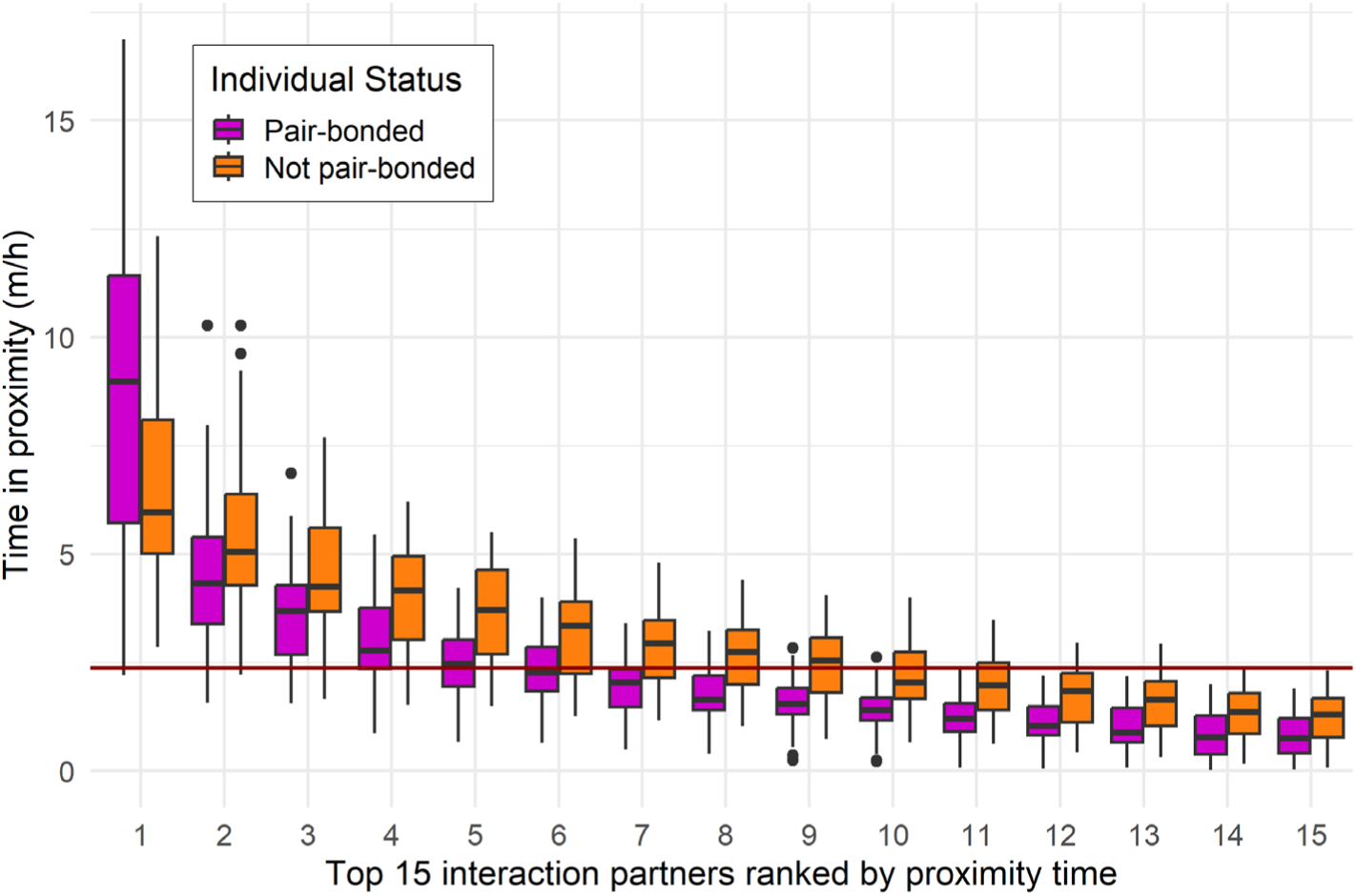
Boxplot showing the proximity rates (minutes per hour a dyad was observed off the nest) with the top 15 interaction partners ranked by time in proximity, plotted for each individual and each period (n = 79 subjects across 4 periods, of which 40 pair-bonded), with pair-bonded and unpaired individuals coloured. Only the first 15 most preferred partners are displayed as group sizes ranged from 16 to 23 individuals. A dark red horizontal line indicates the overall average time in proximity of 2 minutes 16 seconds per hour observed, per observed dyad per period (n = 1432).

To investigate the stability of dyadic bonds, we analysed the correlations of matrices of dyadic bond strengths between all four observation periods using MRQAPs. All four periods were significantly correlated with each other (MRQAP: adj. *P*. < 0.001), with an average correlation coefficient of 0.52 (**Table 1** ). As dyadic proximity rates of previous periods consistently predicted future rates, social bonds were thus relatively stable over time.

**Table 1.**
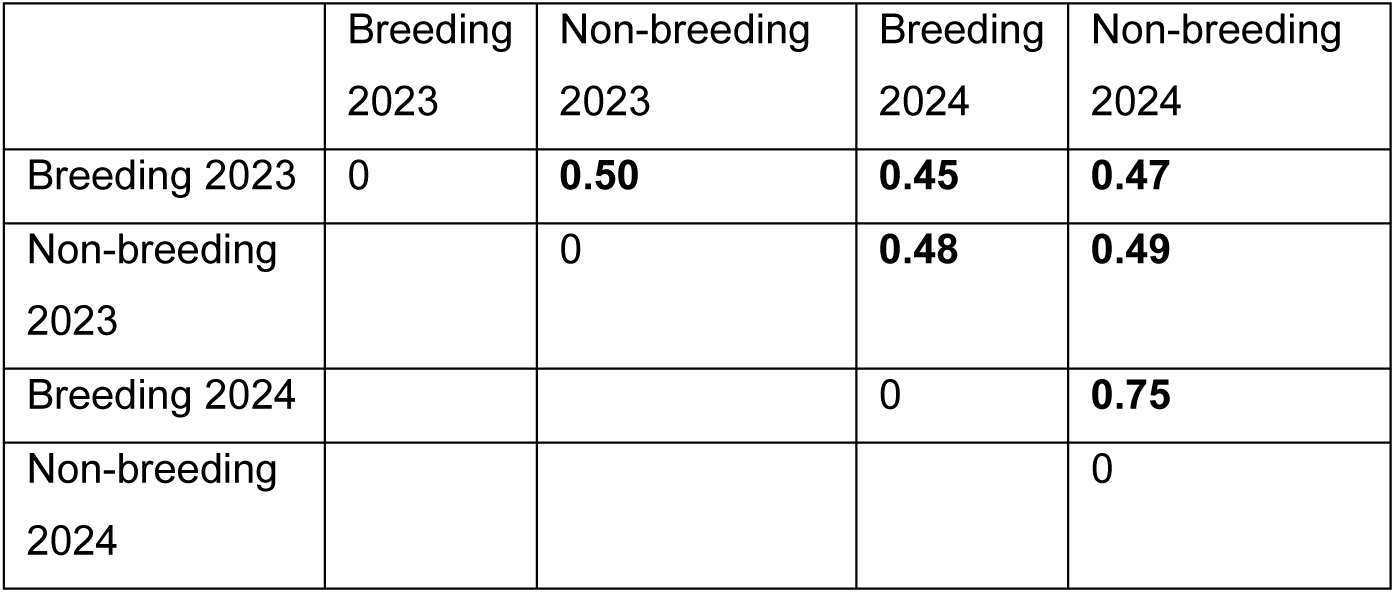
MRQAP correlation coefficient of dyadic proximity rates (m/h) between each two periods. Significant correlations (adj. P< 0.001) are shown in bold.

|  | Breeding<br>2023 | Non-breeding<br>2023 | Breeding<br>2024 | Non-breeding<br>2024 |
| --- | --- | --- | --- | --- |
| Breeding 2023 | 0 | **0.50** | **0.45** | **0.47** |
| Non-breeding<br>2023 |  | 0 | **0.48** | **0.49** |
| Breeding 2024 |  |  | 0 | **0.75** |
| Non-breeding<br>2024 |  |  |  | 0 |

To investigate what shapes social bonds, we examined the dyadic proximity rates during each of the four study periods. Each of the social networks of proximity (m/h) was highly connected (mean density = 0.95 [0.91;0.99], mean transitivity = 0.96 [0.92;0.99]). Any two individuals spent on average about 2 minutes 16 seconds in proximity together per hour observed (range 0 sec. – 16 min. 52 sec. per hour, median 1 min. 44 sec.). While most individuals come into proximity, the wide range in proximity rates, combined with an average of 6.6 strong ties, indicates that social bonds are highly differentiated (**Figure 3**).

To investigate what determines social bond strength in Rüppell’s vultures, we fitted a GLMM on the dyadic proximity rates with age, sex, rank difference, and an interaction of pair-bond and season as predictors, with random effects for dyad identity and period (**model 5, Figure 4**). Sex and rank difference both had no effect on the dyadic bond strength (GLMM: *P* > 0.05, SOM Table S10). Pair-bonded dyads did have substantially higher proximity rates, and this effect varied between the seasons, as we found a significant effect for the interaction between the season and whether a dyad was pair-bonded on dyadic bond strength (GLMM: Breeding season: Pair-bonded: Est. = -1.19 ± 0.25, *P <* 0.001). Thus, we performed post-hoc comparisons to investigate the effects of both pair-bond and season in more detail, averaged over dyadic sex and age (SOM Table S11). Unpaired dyads had weaker bonds than pair-bonded dyads during both the breeding and non-breeding season (Tukey: Est. = -0.87 ± 0.23 and Est. = -2.06 ± 0.19, respectively, adj. *P <* 0.001) (**Figure 5**). Pair-bonded dyads had weaker bonds during the breeding season than outside of the breeding season (Tukey: Est. = 0.99 ± 0.27, adj. *P <* 0.001), while the bonds of unpaired dyads did not differ significantly between the breeding and non-breeding seasons.

**Figure 4.**
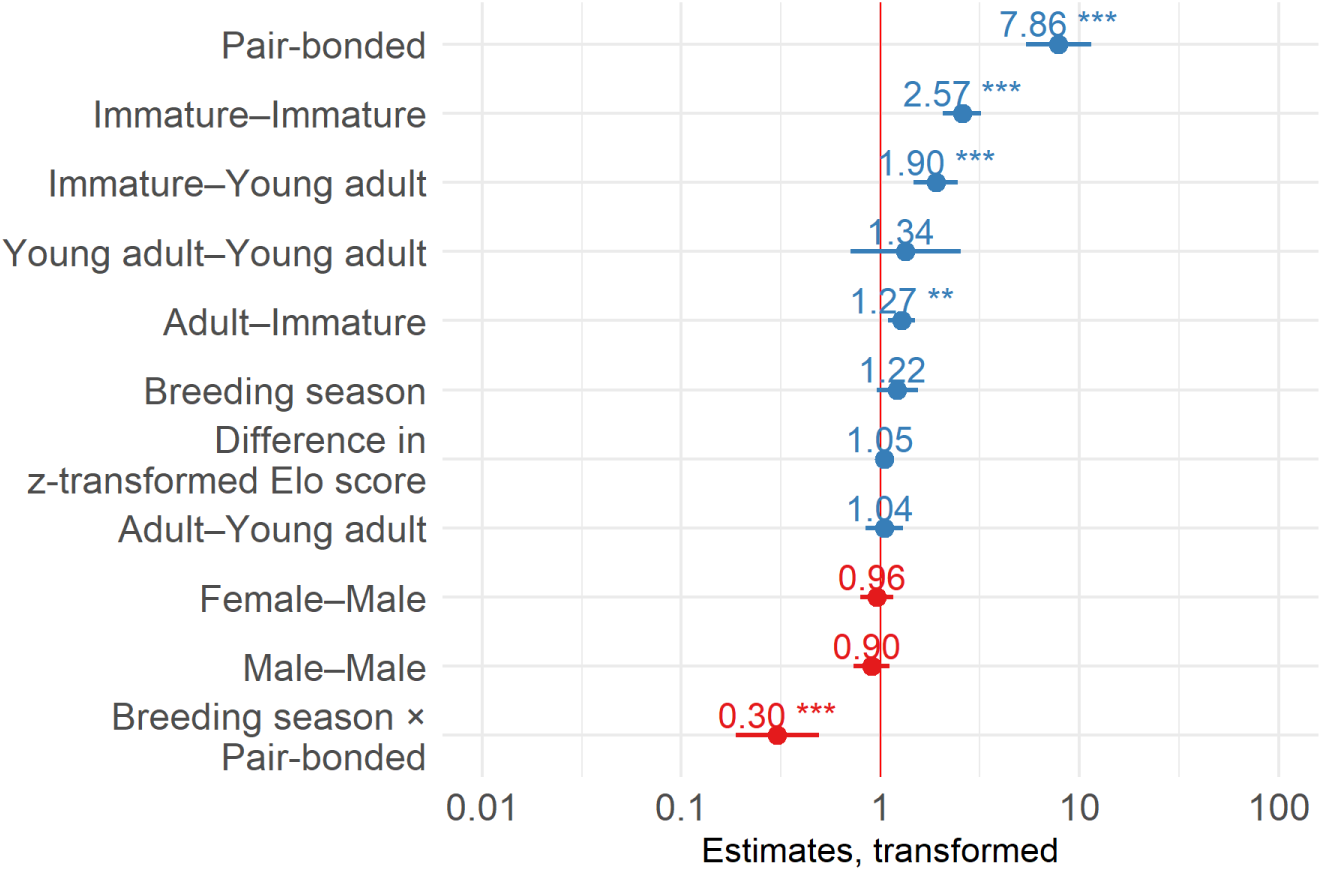
Forest plot of transformed estimates of model 5 predicting dyadic bond strength. Baseline is a non-pair-bonded, adult, female-female dyad during the non-breeding season. Vertical red line indicates no effect. Lines depict 95% confidence interval. Blue horizontal lines and values show positive effects with an estimate > 1 and red depicts negative effects with an estimate < 1. (** P < 0.01, *** P < 0.001)

**Figure 5.**
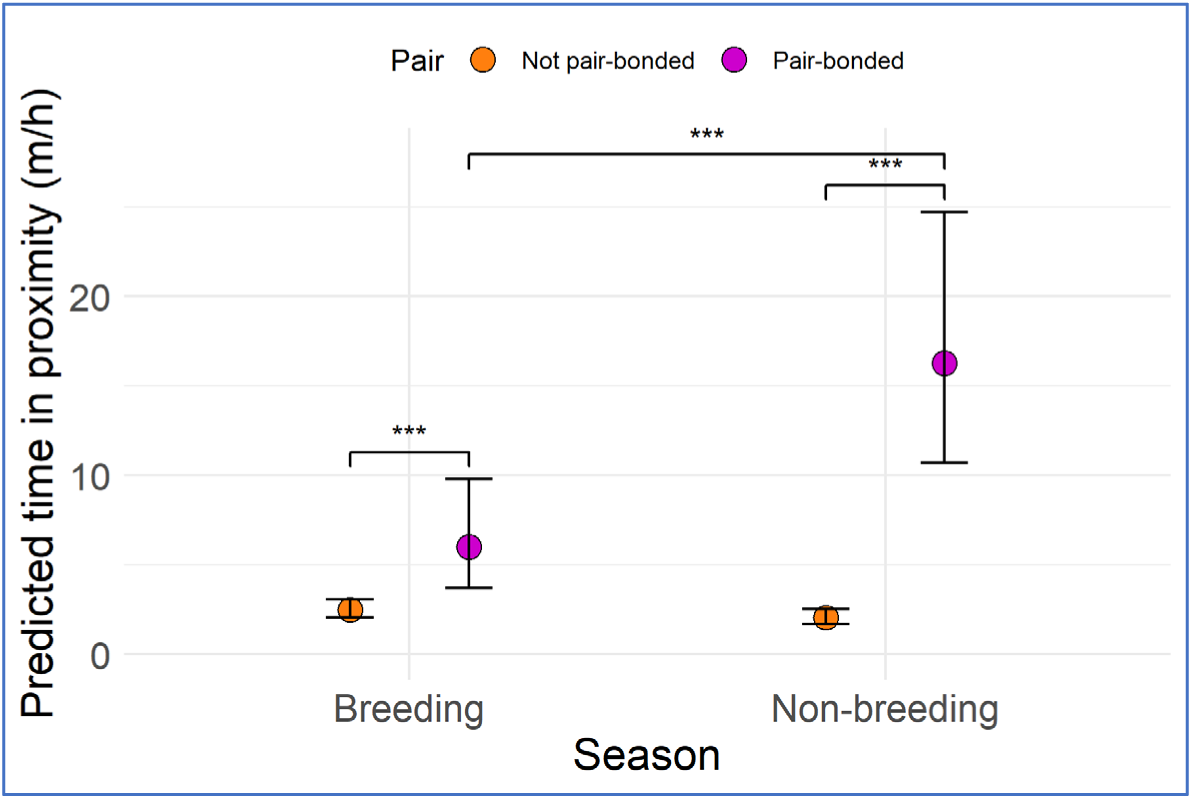
Marginal effect plot for predicted time in proximity (m/h) back-transformed from the log scale plotted against season and pair-bond status, averaged over dyadic sex and age. Error bars reflect standard error. (*** P < 0.001, Tukey adjusted).

In addition to the effects of pair-bond and season, age also had a significant effect on the time a dyad spends in proximity, and thus on social bond strength. A dyad of two immatures, an immature and a young-adult, or an immature and an adult all spent more time together than a baseline of two adults (GLMM: Est. = 0.94 ± 0.11, *P <* 0.001, est. = 0.64 ± 0.13, *P <* 0.001, and est. = 0.24 ± 0.08, *P =* 0.002, respectively). As such, pairwise comparisons were used to investigate the differences between age classes averaged over season, pair-bond status and dyadic sex (Figure 6, SOM Table S11). Dyads involving at least one adult spent 38% to 78% less time together than dyads of at least one immature. This suggests that younger dyads had stronger social bonds than adult dyads.

**Figure 6.**
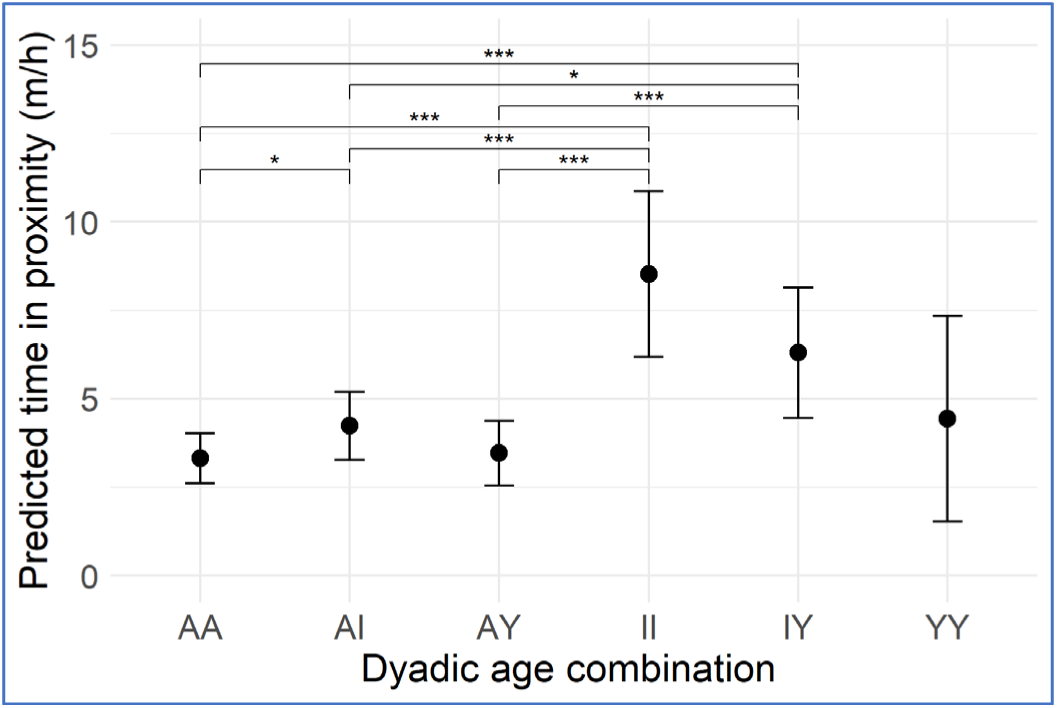
Marginal effects plot of predicted time in proximity (m/h) back-transformed from the log scale plotted against dyadic age combination, averaged over dyadic sex, season and pair-bond status. Error bars reflect standard error. Dyadic age combinations are between adult (A), young adult (Y) and immature (I). (* P < 0.05, ** P < 0.01, *** P < 0.001, Tukey adjusted).

## DISCUSSION

We set out to map the social structure of Rüppell’s vultures by focusing on the nature and patterning of their dominance relationships and affiliative social bonds. Our results revealed that the dominance hierarchy of Rüppell’s vultures was highly stable, as rank reversals were rare, and affiliative bonds were stable as well. However, there were also some seasonal fluctuations in dominance relationships, individual sociality, and dyadic social bond strength. During the breeding season, pair-bonded individuals were more dominant and less social than unpaired individuals, and age, pair-bond status, and season also all influenced the dyadic bond strength. Most notably, pair-bonded dyads had substantially stronger bonds, but maintained strong affiliative bonds outside of this pair-bond too.

The dominance hierarchy we found was intermediately steep and the dominance ranks were highly stable during the course of our study, similar to the hierarchy of another group of adult, captive *Gyps* vultures (7 griffon and 1 Rüppell’s vulture; Sassi et al., 2024). In wild griffon vultures, a strong positive correlation between the number of attacks by an individual and the number of interactions won also suggests a steep dominance hierarchy (Bosè and Sarrazin, 2007), which would imply that individuals engage only when they expect to win. In greylag geese (*Anser anser*), a long-lived bird living in large groups, aggressiveness is a personality trait that predicts dominance rank (Kleindorfer et al., 2024). As such, aggressiveness has been suggested to help maintain stable hierarchies in complex social systems. Therefore, the observed aggressiveness of individual vultures around a carcass could potentially also inform us of their dominance ranks (cf. Boucherie et al., 2022). While the dominance hierarchy of wild *Gyps* vulture populations is as of yet unknown, animal societies are typically structured by some sort of hierarchy (Strauss et al., 2022). Stable dominance hierarchies can form even in large, open societies with highly dynamic social associations such as that of sulphur-crested cockatoos (*Cacatua galerita*) (Penndorf et al., 2026b). Therefore, stable dominance hierarchies could also form in wild, colonially breeding *Gyps* vultures. Studies on other scavenging birds also find stable and steep hierarchies in both wild and captive foraging groups of ravens (Boucherie et al., 2022) and wild Carrion Crows (*C. corone*) (Chiarati et al., 2010). Future work would benefit from studying the dominance hierarchy of Rüppell’s vultures in a wild setting, where the motivation to access a carcass may be higher, group membership may be more variable and other ecological factors might influence the interaction patterns, such as type of carcass, distance to the breeding colony and presence of mammalian scavengers and other vulture species.

We found that Rüppell’s vulture females were more dominant than males. In Egyptian vultures (*Neophron percnopterus*), females are also dominant over males (van Overveld et al., 2018). For birds of prey, it is typical that females are dominant over male partners as they are heavier than males, something referred to as “reversed sexual size dimorphism” (RSD) (see Krüger, 2005; Negro and Galván, 2018). Our finding that female Rüppell’s vultures were more dominant than males therefore fits with the general pattern for birds of prey. However, RSD is believed to be minimal in carrion eating raptors, and female griffon vultures are indeed only marginally larger than males (Xirouchakis and Poulakakis, 2008). The same is likely true for Rüppell’s vultures (Mundy et al., 1992), and while this thus would only give females a slight competitive advantage at most, we nonetheless find a pronounced sex difference in the competitive ability of captive Rüppell’s vultures.

In contrast, a previous study on captive griffon vultures (including one Rüppell’s vulture) finds no effect of sex on dominance (Sassi et al., 2024). In a population of wild-bred and reintroduced griffon vultures in which all individuals are marked, females also do not win a higher proportion of attacks than males, although they do display more frequently than males while feeding (Bosè and Sarrazin, 2007). In our analyses, we combined ritualized interactions and decided aggression, as dominance ranks derived from these two types of interactions are often comparable (Smit, 2024). To test whether the effect of sex on rank stems from differences in threat displays, we compared the number of threats and agonistic interactions of males and females post hoc. Like wild griffon vultures, male Rüppell’s vultures performed 40% fewer threat displays (GLMM: Est = -0.92, SD = 0.25, *P <* 0.001, do note within-group deviation from uniformity for females) but males were also observed to win 42% fewer agonistic interactions than females (GLMM: Est. = -0.88, SD = 0.25, *P <* 0.001). In this captive population of Rüppell’s vultures, females thus displayed more and won more agonistic interactions. As no sex differences were reported in captive griffon vultures and only display rates varied with sex in wild griffons, further investigation of sex effects on dominance rank both in Rüppell’s and griffon vultures is warranted. Moreover, it could be interesting to study the types of agonistic interactions, including undecided aggressions, in more detail to see whether these vary depending on rank differences and to investigate whether agonistic interactions are a strategic investment, as is the case in another social species with a steep hierarchy, the vulturine guineafowl (*Acryllium vulturinum*) (Dehnen et al., 2022).

During the breeding season, pair-bonded Rüppell’s vultures were significantly more dominant than individuals without a pair bond, regardless of whether there was a dependent chick present. In jackdaws (*C. monedula*), which also form long-term pair-bonds and breed colonially, pair-bonded individuals are generally more dominant than unpaired individuals (Kubitza et al., 2015). In greylag geese, paired birds also have higher dominance ranks (Kleindorfer et al., 2024). Studies on juvenile rooks (*C. frugilegus*) and non-breeding aggregations of ravens have also found that individuals with an affiliative or pair-bond are more dominant than those without (Braun and Bugnyar, 2012; Emery et al., 2007). Therefore, pair-bond status having a strong positive effect on dominance only during the breeding season would fit the idea that the social environment of birds is more predictable during the breeding season, as dominance relationships are more clearly defined and based on social roles (Boucherie et al., 2019). Finally, given the stability of these pair-bonds and the dominance hierarchy of the Rüppell’s vultures, it would be interesting to study how this affects their social cognition (cf. SIH; Jolly, 1966; Humphrey, 1976; Dunbar and Shultz, 2007), to investigate what these birds know about their own and others’ social bonds (cf. Massen et al., 2014a) and to test whether they actively act upon this knowledge in a Machiavellian fashion (cf. Massen et al., 2014b, 2025).

We found no effect of age on dominance rank, so immature and young adult Rüppell’s vultures did not have lower ranks than adults. This is similar to another captive, mixed sex group of seven griffon and one Rüppell’s vulture, where age also did not affect rank, although their sample lacked individuals under 4 years of age (Sassi et al., 2024). In contrast to captivity, in wild griffon vultures, adult individuals are considered more dominant than other age classes (Bosè and Sarrazin, 2007) as they initiate more agonistic interactions, and such interactions are often won (Duriez et al., 2012). Younger individuals arrive later at carcasses, when group sizes are larger, and they thus suffer more competition by interference and exploitation, indicating a lower rank. In captivity, young individuals may have easier access to food, resulting in a less pronounced difference in dominance between age classes. Wild studies determine individual age of *Gyps* vultures based on plumage and other physical characteristics (e.g. Duriez et al., 2011; Rodríguez and Elorriaga, 2016). Despite living in captivity, the ages of ten Rüppell’s vultures in our sample are unknown apart from an estimated minimum age. As these are long-lived birds whose sociality also changes with age when tracking individual wild birds (Acácio et al., 2024), it would be informative for future research to test whether the exact age influences dominance ranks in a captive population where ages are known.

With regard to affiliative interactions, during the breeding season, individuals with a pair-bond spent less time in proximity, were less central in the network, and had fewer strong bonds than unpaired individuals. This could be the result of pair-bonded individuals being preoccupied with parental care duties, and therefore becoming more selective in their affiliative bonds. Pair-bonded individuals had fewer strong bonds than unpaired individuals during the breeding season, but it is noteworthy that they still maintained multiple strong bonds. Their affiliative bonds were thus not restricted to (future) reproductive partners, similar to other monogamous scavengers like ravens and rooks (Boucherie et al., 2016; Braun and Bugnyar, 2012; Fraser and Bugnyar, 2010). Captive nonbreeding ravens are selective in their affiliative bonds, forming valuable relationships with few others (Fraser and Bugnyar, 2010). However, where wild, nonbreeding ravens typically focus on just a single bonding partner at a time (Braun and Bugnyar, 2012), we found that both pair-bonded and unpaired captive Rüppell’s vultures maintained multiple strong bonds during 6-month intervals, although it would be valuable to determine the number of strong bonds at a shorter time interval to see to what extent individuals maintain their strong bonds simultaneously.

Although individual sociality and dyadic social bonds were influenced by the breeding and non-breeding season, dyadic affiliative bonds remained stable over time. Even in species where pairs split off from the group during the breeding season, individuals are able to maintain stable social relationships, such as in Sulphur-crested Cockatoos (Penndorf et al., 2026a), and maintain social cohesion, as in vulturine guineafowl (Farine, 2025; Papageorgiou et al., 2019). Colonial *Gyps* vultures may also experience changes in group membership, as the number of griffon vultures on breeding colonies increases during the pre-breeding and incubation period, when numbers on communal roosts decrease (Xirouchakis, 2007). The breeding season of *Gyps* vultures lasts half a year, so they potentially have the opportunity to form and maintain stable bonds both on communal roosts and on breeding colonies. One important note is that the observed stability of Rüppell’s vulture social bonds cannot solely be attributed to the long-term pair-bonds present in the studied population, as the population of up to 23 individuals included five breeding pairs and most of the 253 possible dyads therefore do not share a pair-bond. The stability of these non-pair dyadic bonds we observed is different from nonbreeding ravens in a foraging aggregation, as most nonbreeding ravens do not maintain their affiliative bonds over a two-year period (Braun and Bugnyar, 2012), although it should be noted that the latter study is a wild study, whereas ours is a captive study in which animals are restricted in their choice of partners over the years. Therefore, it would be valuable to track the social bonds of non-breeders and especially immatures over time and investigate what predicts the formation and maintenance of strong ties, particularly in the wild

Multiple factors were predictive of the dyadic bond strength. First, while pair-bonded individuals were less social during the breeding season, their pair-bonds were nonetheless the strongest dyadic bonds. We based social bond strength on proximity while off the nest and, as a result, the time pair-bonded individuals spend in proximity does not reflect attraction to a shared nest. Although we could have missed pair bonded individuals associating on their nest (see Acácio et al., 2024), in the wild, pairs of griffon vultures take turns going on foraging trips while incubating and caring for a chick (Xirouchakis, 2007; Xirouchakis and Mylonas, 2007). Similarly, Rüppell’s vulture breeding pairs may therefore actually spend less time together on the nest during the breeding season. As we found that pair-bonded dyads had stronger bonds outside of the breeding season, this pattern seems to persist even in captivity where food is readily available and the distance back to the nest after feeding is easily traversed.

In addition to season and pair-bond status, age also had an effect on social bond strength as younger individuals had stronger bonds than adult dyads. This fits with the finding that in wild griffon vultures, individuals show a decrease in average bond strength as they age (Acácio et al., 2024). They based bond strength on co-roosting relationships identified using GPS locations, tracking individuals for up to 12 years. We used data collected over two years and correcting for individual identity, enabling us to compare age classes and conclude that younger individuals had stronger bonds than adult individuals. However, as *Gyps* vultures are long-lived birds, we cannot draw meaningful conclusions about changes in sociality as individuals age in Rüppell’s vulture using this dataset alone. This underlines the importance of longitudinal research in such a long-lived species as the Rüppell’s vulture, to test whether the average bond strength of individual birds also decreases as they age.

Co-roosting data has been used to determine bond strength in order to track sociality as griffon vultures age (Acácio et al., 2024), but later work using GPS tracking data has shown that interactions on the ground during the day while feeding and sunbathing contribute to bond strength too (Sharma et al., 2023). While wild studies on griffon vultures often use GPS tracking to determine when and where individuals co-occur (e.g. at night roosts; Acácio et al., 2024), here we used direct observations of social proximity to determine social bond strength of captive Rüppell’s vultures, which could thus give a more detailed (higher resolution) insight into vulture social bond strength. Despite living in captivity, where the connectivity of individuals is likely higher than in an open landscape, we still identified differentiated social bonds that were shaped by individual characteristics, pair-bond status, dominance relationships, and season. Although a single captive population of Rüppell’s vultures is likely not representative of all other *Gyps* vulture populations, both wild and captive, studying a captive population in detail does allow us to gain a greater understanding of the potential complexity of vulture social life and the potential diversity in their dominance and affiliative relationships.

We aimed to investigate the dominance and affiliative relationships among Rüppell’s vultures, to study the potential social complexity experienced by these birds. We conclude that multiple forces shape the dominance hierarchy and affiliative network of a captive group of Rüppell’s vultures. These detailed insights into what governs vulture social interactions allow us to better quantify the extent of relationship differentiation and of social uncertainty (Hobson et al., 2019; Kappeler et al., 2019). The effects of individual characteristics on dominance relationships, individual sociality and dyadic bond strength suggest that they form different types of social bonds apart from the pair-bond, and social bonds are thus differentiated, adding to the complexity of this system. In corvids and parrots, individuals also form multiple individualized relationships and use social information in decision making processes, markers of their “complex” social structure (Boucherie et al., 2019; Penndorf et al., 2026b, 2025). Some social uncertainty exists when it comes to social bonds, as both dominance relationships and affiliative social bonds show seasonal fluctuations. This seasonal effect is in line with the notion that the social environment of birds is more predictable during the breeding than the non-breeding season (Boucherie et al., 2019). Combined, the current metrics would suggest that captive Rüppell’s vultures live in a complex social environment and share aspects of their social structure with corvid and parrot species. While the cognitive abilities of vultures are yet to be studied in detail, following the SIH, the complex social environment of *Gyps* vultures may have resulted in the evolution of potentially elaborate cognitive abilities, just like it has in corvids and parrots.

Here, we investigated dominance relationships and social bonds to identify potential sources of social complexity in the social structure of Rüppell’s vultures, including relationship differentiation and social uncertainty. While a lot of different metrics can be involved in estimating social complexity, a recent meta-analysis across different taxa finds that support for the SIH is not predicted by which measure of sociality is used when comparing broad metrics on the social system, group size or individual-level social interactions (Speechley et al., 2024). On the other hand, the nuances of individual-level social complexity could also influence individual cognitive performance. For example, sex-specific differences in social interactions have been shown to influence the performance of budgerigars (*Melopsittacus undulatus*) (Tang et al., 2026) and cooperatively breeding Australian magpies (*Cracticus tibicen dorsalis*) that live in larger groups have increased cognitive performance (Ashton et al., 2018). Therefore, mapping the social structure of more populations of *Gyps* vultures as well as less social vulture species in detail would allow for a detailed comparison of social complexity and reveal whether a complex social environment may have been a selection pressure on the evolution of intelligence in general, and in this promising new taxonomic group for cognition studies, specifically.

## Data Availability Statement

*The data and code for reproducing our analysis will be provided after completing the peer review process*.

## Acknowledgements

We thank the members of the Vulture Cognition Lab at Utrecht University and especially Connor Proudfoot for fruitful discussions, and Maarten Vis of Royal Rotterdam Zoological & Botanical Gardens, The Netherlands, for his insights as coordinator of the EAZA Ex-situ Program for the Rüppell’s vulture. Special thanks go to Joost Jammers from Avifauna Bird Park for allowing us to study the Rüppell’s vultures in their care. We also thank Dennis Appels for his help during the start of this research. Finally, we are very grateful to the dedicated caretakers at Avifauna for their ongoing support and excellent care of the vultures.

Financial support for this study was provided by the Ministry of Education, Culture and Science of the Netherlands through a starters and incentive grant (#U2023-169) to J.J.M.M.

## References

Acácio, M., Gahm, K., Anglister, N., Vaadia, G., Hatzofe, O., Harel, R., Efrat, R., Nathan, R., Pinter-Wollman, N., Spiegel, O., 2024. Behavioral plasticity shapes population aging patterns in a long-lived avian scavenger. Proc. Natl. Acad. Sci. 121, e2407298121. 10.1073/pnas.2407298121

Allen, C., 2017. On (not) defining cognition. Synthese 194, 4233–4249. 10.1007/s11229-017-1454-4

Ashton, B.J., Ridley, A.R., Edwards, E.K., Thornton, A., 2018. Cognitive performance is linked to group size and affects fitness in Australian magpies. Nature 554, 364–367. 10.1038/nature25503

Balasubramaniam, K.N., Dittmar, K., Berman, C.M., Butovskaya, M., Cooper, M.A., Majolo, B., Ogawa, H., Schino, G., Thierry, B., de Waal, F.B.M., 2012. Hierarchical steepness and phylogenetic models: phylogenetic signals in *Macaca*. Anim. Behav. 83, 1207–1218. 10.1016/j.anbehav.2012.02.012

Bildstein, K.L., 2022. Vultures of the World: Essential Ecology and Conservation. Cornell University Press.

Borgeaud, C., Sosa, S., Sueur, C., Bshary, R., 2017. The influence of demographic variation on social network stability in wild vervet monkeys. Anim. Behav. 134, 155–165. 10.1016/j.anbehav.2017.09.028

Bosè, M., Duriez, O., Sarrazin, F., 2012. Intra-specific competition in foraging griffon vultures Gyps fulvus: 1. Dynamics of group feeding. Bird Study 59, 182–192. 10.1080/00063657.2012.658639

Bosè, M., Sarrazin, F., 2007. Competitive behaviour and feeding rate in a reintroduced population of griffon vultures Gyps fulvus. Ibis 149, 490–501. 10.1111/j.1474-919X.2007.00674.x

Boucherie, P.H., Gallego-Abenza, M., Massen, J.J.M., Bugnyar, T., 2022. Dominance in a socially dynamic setting: hierarchical structure and conflict dynamics in ravens’ foraging groups. Philos. Trans. R. Soc. B Biol. Sci. 377, 20200446. 10.1098/rstb.2020.0446

Boucherie, P.H., Loretto, M.-C., Massen, J.J.M., Bugnyar, T., 2019. What constitutes “social complexity” and “social intelligence” in birds? Lessons from ravens. Behav. Ecol. Sociobiol. 73, 12. 10.1007/s00265-018-2607-2

Boucherie, P.H., Mariette, M.M., Bret, C., Dufour, V., 2016. Bonding beyond the pair in a monogamous bird: impact on social structure in adult rooks (Corvus frugilegus). Behaviour 153, 897–925. 10.1163/1568539X-00003372

Bräuer, J., Hanus, D., Pika, S., Gray, R., Uomini, N., 2020. Old and New Approaches to Animal Cognition: There Is Not “One Cognition.” J. Intell. 8, 28. 10.3390/jintelligence8030028

Braun, A., Bugnyar, T., 2012. Social bonds and rank acquisition in raven nonbreeder aggregations. Anim. Behav. 84, 1507–1515. 10.1016/j.anbehav.2012.09.024

Brent, L.J.N., Semple, S., Dubuc, C., Heistermann, M., MacLarnon, A., 2011. Social capital and physiological stress levels in free-ranging adult female rhesus macaques. Physiol. Behav. 102, 76–83. 10.1016/j.physbeh.2010.09.022

Brooks, M., Bolker, B., Kristensen, K., Maechler, M., Magnusson, A., McGillycuddy, M., Skaug, H., Nielsen, A., Berg, C., Bentham, K. van, Sadat, N., Lüdecke, D., Lenth, R., O’Brien, J., Geyer, C.J., Jagan, M., Wiernik, B., Stouffer, D.B., Agronah, M., Akdur, H.T.K., Bové, D.S., Krieger, N., Williams, C., 2026. glmmTMB: Generalized Linear Mixed Models using Template Model Builder.

Bugnyar, T., 2024. Why are ravens smart? Exploring the social intelligence hypothesis. J. Ornithol. 165, 15–26. 10.1007/s10336-023-02111-6

Byrne, R.W., Whiten, A., 1988. Machiavellian intelligence: social expertise and the evolution of intellect in monkeys, apes, and humans. Clarendon Press, Oxford.

Cheney, D.L., Seyfarth, R.M., 2007. Baboon metaphysics: the evolution of a social mind. University of Chicago Press, Chicago.

Chiarati, E., Canestrari, D., Vera, R., Marcos, J.M., Baglione, V., 2010. Linear and Stable Dominance Hierarchies in Cooperative Carrion Crows. Ethology 116, 346–356. 10.1111/j.1439-0310.2010.01741.x

Cortés-Avizanda, A., Jovani, R., Donázar, J.A., Grimm, V., 2014. Bird sky networks: How do avian scavengers use social information to find carrion? Ecology 95, 1799–1808. 10.1890/13-0574.1

Damas, J., Corbo, M., Kim, J., Turner-Maier, J., Farré, M., Larkin, D.M., Ryder, O.A., Steiner, C., Houck, M.L., Hall, S., Shiue, L., Thomas, S., Swale, T., Daly, M., Korlach, J., Uliano-Silva, M., Mazzoni, C.J., Birren, B.W., Genereux, D.P., Johnson, J., Lindblad-Toh, K., Karlsson, E.K., Nweeia, M.T., Johnson, R.N., Zoonomia Consortium, Lewin, H.A., 2022. Evolution of the ancestral mammalian karyotype and syntenic regions. Proc. Natl. Acad. Sci. 119, e2209139119. 10.1073/pnas.2209139119

de Vries, H., Stevens, J.M.G., Vervaecke, H., 2006. Measuring and testing the steepness of dominance hierarchies. Anim. Behav. 71, 585–592. 10.1016/j.anbehav.2005.05.015

Deakin, J.E., Ezaz, T., 2014. Tracing the evolution of amniote chromosomes. Chromosoma 123, 201. 10.1007/s00412-014-0456-y

Dehnen, T., Papageorgiou, D., Nyaguthii, B., Cherono, W., Penndorf, J., Boogert, N.J., Farine, D.R., 2022. Costs dictate strategic investment in dominance interactions. Philos. Trans. R. Soc. B Biol. Sci. 377, 20200447. 10.1098/rstb.2020.0447

Dekker, D., Krackhardt, D., Snijders, T.A.B., 2007. Sensitivity of MRQAP Tests to Collinearity and Autocorrelation Conditions. Psychometrika 72, 563–581. 10.1007/s11336-007-9016-1

Dunbar, R.I.M., Shultz, S., 2007. Evolution in the Social Brain. Science 317, 1344–1347. 10.1126/science.1145463

Duriez, O., Eliotout, Bertrand, and Sarrazin, F., 2011. Age identification of Eurasian griffon vultures Gyps fulvus in the field. Ringing Migr. 26, 24–30. 10.1080/03078698.2011.585912

Duriez, O., Herman, S., Sarrazin, F., 2012. Intra-specific competition in foraging griffon vultures Gyps fulvus: 2. The influence of supplementary feeding management. Bird Study 59, 193–206. 10.1080/00063657.2012.658640

Emery, N.J., 2005. Cognitive ornithology: the evolution of avian intelligence. Philos. Trans. R. Soc. B Biol. Sci. 361, 23–43. 10.1098/rstb.2005.1736

Emery, N.J., Seed, A.M., von Bayern, A.M.P., Clayton, N.S., 2007. Cognitive adaptations of social bonding in birds. Philos. Trans. R. Soc. B Biol. Sci. 362, 489–505. 10.1098/rstb.2006.1991

European Association of Zoos and Aquaria, 2024. EAZA Standards for Welfare, Accommodation and Management of Animals in Zoos and Aquariums.

Farine, D.R., 2025. The multifaceted societies of vulturine guineafowl. Anim. Behav. 229, 123333. 10.1016/j.anbehav.2025.123333

Fraser, O.N., Bugnyar, T., 2010. The quality of social relationships in ravens. Anim. Behav. 79, 927–933. 10.1016/j.anbehav.2010.01.008

Freeberg, T.M., Dunbar, R.I.M., Ord, T.J., 2012. Social complexity as a proximate and ultimate factor in communicative complexity. Philos. Trans. R. Soc. B Biol. Sci. 367, 1785–1801. 10.1098/rstb.2011.0213

Friard, O., Gamba, M., 2016. BORIS: a free, versatile open-source event-logging software for video/audio coding and live observations. Methods Ecol. Evol. 7, 1325–1330. 10.1111/2041-210X.12584

Güntürkün, O., Ströckens, F., Scarf, D., Colombo, M., 2017. Apes, feathered apes, and pigeons: differences and similarities. Curr. Opin. Behav. Sci., Comparative cognition 16, 35–40. 10.1016/j.cobeha.2017.03.003

Harel, R., Spiegel, O., Getz, W.M., Nathan, R., 2017. Social foraging and individual consistency in following behaviour: testing the information centre hypothesis in free-ranging vultures. Proc. R. Soc. B Biol. Sci. 284, 20162654. 10.1098/rspb.2016.2654

Hartig, F., Lohse, L., leite, M. de S., 2024. DHARMa: Residual Diagnostics for Hierarchical (Multi-Level / Mixed) Regression Models.

Hirschauer, M.T., Wolter, K., 2017. High occurrence of extra-pair partnerships and homosexuality in a captive Cape vulture Gyps coprotheres colony§. Ostrich 88, 173–176. 10.2989/00306525.2017.1304461

Hobson, E.A., Ferdinand, V., Kolchinsky, A., Garland, J., 2019. Rethinking animal social complexity measures with the help of complex systems concepts. Anim. Behav. 155, 287–296. 10.1016/j.anbehav.2019.05.016

Humphrey, N.K., 1976. The social function of intellect. Grow. Points Ethol. 303–317.

Jarvis, E.D., Mirarab, S., Aberer, A.J., Li, B., Houde, P., Li, C., Ho, S.Y.W., Faircloth, B.C., Nabholz, B., Howard, J.T., Suh, A., Weber, C.C., da Fonseca, R.R., Li, J., Zhang, F., Li, H., Zhou, L., Narula, N., Liu, L., Ganapathy, G., Boussau, B., Bayzid, Md.S., Zavidovych, V., Subramanian, S., Gabaldón, T., Capella-Gutiérrez, S., Huerta-Cepas, J., Rekepalli, B., Munch, K., Schierup, M., Lindow, B., Warren, W.C., Ray, D., Green, R.E., Bruford, M.W., Zhan, X., Dixon, A., Li, S., Li, N., Huang, Y., Derryberry, E.P., Bertelsen, M.F., Sheldon, F.H., Brumfield, R.T., Mello, C.V., Lovell, P.V., Wirthlin, M., Schneider, M.P.C., Prosdocimi, F., Samaniego, J.A., Velazquez, A.M.V., Alfaro-Núñez, A., Campos, P.F., Petersen, B., Sicheritz-Ponten, T., Pas, A., Bailey, T., Scofield, P., Bunce, M., Lambert, D.M., Zhou, Q., Perelman, P., Driskell, A.C., Shapiro, B., Xiong, Z., Zeng, Y., Liu, S., Li, Z., Liu, B., Wu, K., Xiao, J., Yinqi, X., Zheng, Q., Zhang, Y., Yang, H., Wang, J., Smeds, L., Rheindt, F.E., Braun, M., Fjeldsa, J., Orlando, L., Barker, F.K., Jønsson, K.A., Johnson, W., Koepfli, K.-P., O’Brien, S., Haussler, D., Ryder, O.A., Rahbek, C., Willerslev, E., Graves, G.R., Glenn, T.C., McCormack, J., Burt, D., Ellegren, H., Alström, P., Edwards, S.V., Stamatakis, A., Mindell, D.P., Cracraft, J., Braun, E.L., Warnow, T., Jun, W., Gilbert, M.T.P., Zhang, G., 2014. Whole-genome analyses resolve early branches in the tree of life of modern birds. Science 346, 1320–1331. 10.1126/science.1253451

Jolly, A., 1966. Lemur Social Behavior and Primate Intelligence. Science 153, 501–506. 10.1126/science.153.3735.501

Kane, A., Monadjem, A., Aschenborn, H.K.O., Bildstein, K., Botha, A., Bracebridge, C., Buechley, E.R., Buij, R., Davies, J.P., Diekmann, M., Downs, C.T., Farwig, N., Galligan, T., Kaltenecker, G., Kelly, C., Kemp, R., Kolberg, H., MacKenzie, M.L., Mendelsohn, J., Mgumba, M., Nathan, R., Nicholas, A., Ogada, D., Pfeiffer, M.B., Phipps, W.L., Pretorius, M.D., Rösner, S., Schabo, D.G., Shatumbu, G.L., Spiegel, O., Thompson, L.J., Venter, J.A., Virani, M., Wolter, K., Kendall, C.J., 2022. Understanding continent-wide variation in vulture ranging behavior to assess feasibility of vulture Safe Zones in Africa: Challenges and possibilities. Biol. Conserv. 268, 109516. 10.1016/j.biocon.2022.109516

Kappeler, P.M., 2019. A framework for studying social complexity. Behav. Ecol. Sociobiol. 73, 13. 10.1007/s00265-018-2601-8

Kappeler, P.M., Clutton-Brock, T., Shultz, S., Lukas, D., 2019. Social complexity: patterns, processes, and evolution. Behav. Ecol. Sociobiol. 73, 5. 10.1007/s00265-018-2613-4

Kleindorfer, S., Krupka, M.A., Katsis, A.C., Frigerio, D., Common, L.K., 2024. Aggressiveness predicts dominance rank in greylag geese: mirror tests and agonistic interactions. R. Soc. Open Sci. 11, 231686. 10.1098/rsos.231686

Koo, T.K., Li, M.Y., 2016. A Guideline of Selecting and Reporting Intraclass Correlation Coefficients for Reliability Research. J. Chiropr. Med. 15, 155–163. 10.1016/j.jcm.2016.02.012

Krackhardt, D., 1988. Predicting with networks: Nonparametric multiple regression analysis of dyadic data. Soc. Netw. 10, 359–381. 10.1016/0378-8733(88)90004-4

Krüger, O., 2005. The Evolution of Reversed Sexual Size Dimorphism in Hawks, Falcons and Owls: A Comparative Study. Evol. Ecol. 19, 467–486. 10.1007/s10682-005-0293-9

Kubitza, R.J., Bugnyar, T., Schwab, C., 2015. Pair bond characteristics and maintenance in free-flying jackdaws Corvus monedula: effects of social context and season. J. Avian Biol. 46, 206–215. 10.1111/jav.00508

Lambert, M.L., Jacobs, I., Osvath, M., Bayern, A.M.P. von, 2019. Birds of a feather? Parrot and corvid cognition compared. Behaviour 156, 505–594. 10.1163/1568539X-00003527

Lenth, R.V., Piaskowski, J., Banfai, B., Bolker, B., Buerkner, P., Giné-Vázquez, I., Hervé, M., Jung, M., Love, J., Miguez, F., Riebl, H., Singmann, H., 2026. emmeans: Estimated Marginal Means, aka Least-Squares Means.

Massen, J.J.M., 2021. Studying the Evolution of Cooperation and Prosociality in Birds. Proc. Annu. Meet. Cogn. Sci. Soc. 43.

Massen, J.J.M., Boucherie, P.H., Bugnyar, T., 2025. Third-party interventions of common ravens, *Corvus corax*. Anim. Behav. 219, 123004. 10.1016/j.anbehav.2024.10.018

Massen, J.J.M., Pašukonis, A., Schmidt, J., Bugnyar, T., 2014a. Ravens notice dominance reversals among conspecifics within and outside their social group. Nat. Commun. 5, 3679. 10.1038/ncomms4679

Massen, J.J.M., Sterck, E., Vos, H. de, 2010. Close social associations in animals and humans: functions and mechanisms of friendship. Behaviour 147, 1379–1412. 10.1163/000579510X528224

Massen, J.J.M., Sterck, E.H.M., 2013. Stability and Durability of Intra- and Intersex Social Bonds of Captive Rhesus Macaques (Macaca mulatta). Int. J. Primatol. 34, 770–791. 10.1007/s10764-013-9695-7

Massen, J.J.M., Szipl, G., Spreafico, M., Bugnyar, T., 2014b. Ravens Intervene in Others’ Bonding Attempts. Curr. Biol. 24, 2733–2736. 10.1016/j.cub.2014.09.073

McDonald, D.B., Shizuka, D., 2013. Comparative transitive and temporal orderliness in dominance networks. Behav. Ecol. 24, 511–520. 10.1093/beheco/ars192

McFarland, R., Murphy, D., Lusseau, D., Henzi, S.P., Parker, J.L., Pollet, T.V., Barrett, L., 2017. The ‘strength of weak ties’ among female baboons: fitness-related benefits of social bonds. Anim. Behav. 126, 101–106. 10.1016/j.anbehav.2017.02.002

Milton, K., 1981. Distribution Patterns of Tropical Plant Foods as an Evolutionary Stimulus to Primate Mental Development. Am. Anthropol. 83, 534–548. 10.1525/aa.1981.833.02a00020

Moll, H., Tomasello, M., 2007. Cooperation and human cognition: the Vygotskian intelligence hypothesis. Philos. Trans. R. Soc. B Biol. Sci. 362, 639–648. 10.1098/rstb.2006.2000

Mundy, P., Butchart, D., Ledger, J., Piper, S.E., 1992. The vultures of Africa.

Negro, J.J., Galván, I., 2018. Behavioural Ecology of Raptors, in: Sarasola, J.H., Grande, J.M., Negro, J.J. (Eds.), Birds of Prey: Biology and Conservation in the XXI Century. Springer International Publishing, Cham, pp. 33–62. 10.1007/978-3-319-73745-4_2

Neumann, C., 2023. EloSteepness: Bayesian Dominance Hierarchy Steepness via Elo Rating and David’s Scores.

Neumann, C., Duboscq, J., Dubuc, C., Ginting, A., Irwan, A.M., Agil, M., Widdig, A., Engelhardt, A., 2011. Assessing dominance hierarchies: validation and advantages of progressive evaluation with Elo-rating. Anim. Behav. 82, 911–921. 10.1016/j.anbehav.2011.07.016

Neumann, C., Fischer, J., 2023. Extending Bayesian Elo-rating to quantify the steepness of dominance hierarchies. Methods Ecol. Evol. 14, 669–682. 10.1111/2041-210X.14021

Neumann, C., Kulik, L., 2024. Animal Dominance Hierarchies by Elo Rating.

Papageorgiou, D., Christensen, C., Gall, G.E.C., Klarevas-Irby, J.A., Nyaguthii, B., Couzin, I.D., Farine, D.R., 2019. The multilevel society of a small-brained bird. Curr. Biol. 29, R1120–R1121. 10.1016/j.cub.2019.09.072

Parker, S.T., Gibson, K.R., 1977. Object manipulation, tool use and sensorimotor intelligence as feeding adaptations in cebus monkeys and great apes. J. Hum. Evol. 6, 623–641. 10.1016/S0047-2484(77)80135-8

Peckre, L., Kappeler, P.M., Fichtel, C., 2019. Clarifying and expanding the social complexity hypothesis for communicative complexity. Behav. Ecol. Sociobiol. 73, 11. 10.1007/s00265-018-2605-4

Penndorf, J., Csak, A., Aplin, L., 2026a. Sex-dependent carry-over between winter social associations and breeding proximity in a communally roosting wild parrot. Biol. Lett. 22, 20250832. 10.1098/rsbl.2025.0832

Penndorf, J., Farine, D., Martin, J., Aplin, L., 2026b. Long-term stability of dominance hierarchies in a wild parrot with fission–fusion dynamics. Anim. Behav. 233, 123478. 10.1016/j.anbehav.2026.123478

Penndorf, J., Farine, D.R., Martin, J.M., Aplin, L.M., 2025. Parrot politics: social decision-making in wild parrots relies on both individual recognition and intrinsic markers. R. Soc. Open Sci. 12, 241542. 10.1098/rsos.241542

Pika, S., Sima, M.J., Blum, C.R., Herrmann, E., Mundry, R., 2020. Ravens parallel great apes in physical and social cognitive skills. Sci. Rep. 10, 20617. 10.1038/s41598-020-77060-8

Posit team, 2023. RStudio: Integrated Development Environment for R.

Proudfoot, C., Brons, C.N., Vis, M.W.A., Schel, A., van Dijk, E.S.J., Massen, J.J.M., in press. The Complex, Graded Vocal Repertoire of Rüppell’s vultures Reflects Social-Cognitive Coevolution. PLOS ONE. 10.2139/ssrn.5732416

Prum, R.O., Berv, J.S., Dornburg, A., Field, D.J., Townsend, J.P., Lemmon, E.M., Lemmon, A.R., 2015. A comprehensive phylogeny of birds (Aves) using targeted next-generation DNA sequencing. Nature 526, 569–573. 10.1038/nature15697

R Core Team, 2025. A Language and Environment for Statistical Computing.

Rodríguez, G., Elorriaga, J., 2016. Identification of Rüppell’s vulture and White-backed vulture and vagrancy in the WP. Dutch Bird. 38, 349–375.

Romani, F., Ramella Levis, E., Posillico, M., Opramolla, G., Pavan, G., 2022. Vocal repertoire of the Eurasian griffon vulture (Gyps fulvus) in the central Apennines: a baseline assessment. Bioacoustics 31, 283–312. 10.1080/09524622.2021.1925591

Rosati, A.G., 2017. Foraging Cognition: Reviving the Ecological Intelligence Hypothesis. Trends Cogn. Sci. 21, 691–702. 10.1016/j.tics.2017.05.011

Sánchez-Tójar, A., Schroeder, J., Farine, D.R., 2018. A practical guide for inferring reliable dominance hierarchies and estimating their uncertainty. J. Anim. Ecol. 87, 594–608. 10.1111/1365-2656.12776

Sassi, Y., Nouzières, B., Scacco, M., Tremblay, Y., Duriez, O., Robira, B., 2024. The use of social information in vulture flight decisions. Proc. R. Soc. B Biol. Sci. 291, 20231729. 10.1098/rspb.2023.1729

Schülke, O., Anzà, S., Crockford, C., De Moor, D., Deschner, T., Fichtel, C., Gogarten, J.F., Kappeler, P.M., Manin, V., Müller-Klein, N., Prox, L., Sadoughi, B., Touitou, S., Wittig, R.M., Ostner, J., 2022. Quantifying within-group variation in sociality—covariation among metrics and patterns across primate groups and species. Behav. Ecol. Sociobiol. 76, 50. 10.1007/s00265-022-03133-5

Sharma, N., Anglister, N., Spiegel, O., Pinter-Wollman, N., 2023. Social situations differ in their contribution to population-level social structure in griffon vultures. Ecol. Evol. 13, e10139. 10.1002/ece3.10139

Smit, N., 2024. Hierarchies inferred from different agonistic behaviours are not always comparable. J. Anim. Ecol. 93, 1947–1959. 10.1111/1365-2656.14203

Smit, N., Robbins, M.M., 2025. Female gorillas form highly stable dominance relationships. Biol. Lett. 21, 20240556. 10.1098/rsbl.2024.0556

Sol, D., Duncan, R.P., Blackburn, T.M., Cassey, P., Lefebvre, L., 2005. Big brains, enhanced cognition, and response of birds to novel environments. Proc. Natl. Acad. Sci. 102, 5460–5465. 10.1073/pnas.0408145102

Speechley, E.M., Ashton, B.J., Foo, Y.Z., Simmons, L.W., Ridley, A.R., 2024. Meta-analyses reveal support for the Social Intelligence Hypothesis. Biol. Rev. 99, 1889–1908. 10.1111/brv.13103

Stiller, J., Feng, S., Chowdhury, A.-A., Rivas-González, I., Duchêne, D.A., Fang, Q., Deng, Y., Kozlov, A., Stamatakis, A., Claramunt, S., Nguyen, J.M.T., Ho, S.Y.W., Faircloth, B.C., Haag, J., Houde, P., Cracraft, J., Balaban, M., Mai, U., Chen, G., Gao, R., Zhou, C., Xie, Y., Huang, Z., Cao, Z., Yan, Z., Ogilvie, H.A., Nakhleh, L., Lindow, B., Morel, B., Fjeldså, J., Hosner, P.A., da Fonseca, R.R., Petersen, B., Tobias, J.A., Székely, T., Kennedy, J.D., Reeve, A.H., Liker, A., Stervander, M., Antunes, A., Tietze, D.T., Bertelsen, M.F., Lei, F., Rahbek, C., Graves, G.R., Schierup, M.H., Warnow, T., Braun, E.L., Gilbert, M.T.P., Jarvis, E.D., Mirarab, S., Zhang, G., 2024. Complexity of avian evolution revealed by family-level genomes. Nature 629, 851–860. 10.1038/s41586-024-07323-1

Strauss, E.D., Curley, J.P., Shizuka, D., Hobson, E.A., 2022. The centennial of the pecking order: current state and future prospects for the study of dominance hierarchies. Philos. Trans. R. Soc. B Biol. Sci. 377, 20200432. 10.1098/rstb.2020.0432

Tang, Z., Wu, X., Zhang, Y., Sun, Y., Lyu, N., 2026. Sex-specific effects of social complexity and competitive interactions on cognitive performance in Budgerigars. Avian Res. 17, 100343. 10.1016/j.avrs.2026.100343

Tibbetts, E.A., Pardo-Sanchez, J., Weise, C., 2022. The establishment and maintenance of dominance hierarchies. Philos. Trans. R. Soc. B Biol. Sci. 377, 20200450. 10.1098/rstb.2020.0450

Van Horik, J.O., Clayton, N.S., Emery, N.J., 2012. Convergent evolution of cognition in corvids, apes and other animals. Oxf. Handb. Comp. Evol. Psychol. 80–101.

van Overveld, T., García-Alfonso, M., Dingemanse, N.J., Bouten, W., Gangoso, L., de la Riva, M., Serrano, D., Donázar, J.A., 2018. Food predictability and social status drive individual resource specializations in a territorial vulture. Sci. Rep. 8, 15155. 10.1038/s41598-018-33564-y

van Overveld, T., Sol, D., Blanco, G., Margalida, A., de la Riva, M., Donázar, J.A., 2022. Vultures as an overlooked model in cognitive ecology. Anim. Cogn. 25, 495–507. 10.1007/s10071-021-01585-2

Weiß, B.M., Kehmeier, S., Schloegl, C., 2010. Transitive inference in free-living greylag geese, *Anser anser*. Anim. Behav. 79, 1277–1283. 10.1016/j.anbehav.2010.02.029

Wink, M., Starikov, I.J., Schulze-Hagen, K., Negro, J.J., Margalida, A., 2026. What, if anything, is a vulture? Phylogeny and trait evolution in obligate avian scavengers. J. Ornithol. 10.1007/s10336-025-02358-1

Xirouchakis, S.M., 2007. Seasonal and daily activity pattern in griffon vulture (Gyps fulvus) colonies on the island of Crete (Greece). Ornis Fenn. 84, 39.

Xirouchakis, S.M., Mylonas, M., 2007. Breeding behaviour and parental care in the griffon vulture Gyps fulvus on the island of Crete (Greece). Ethol. Ecol. Evol. 19, 1–26. 10.1080/08927014.2007.9522578

Xirouchakis, S.M., Poulakakis, N., 2008. Biometrics, Sexual Dimorphism and Gender Determination of griffon vultures Gyps fulvus from Crete. Ardea 96, 91–98. 10.5253/078.096.0110

Yadav Ruby, A.K., Kanaujia, A., 2018. A Review on Aerial Display, Nest site selection, Nest Defense, Courtship display, Copulation, Parental care: key events of vultures during Breeding, threats and Management. Int J Life Sci. 6, 586–604.

