## Supplements for "The social structure of a captive group of Rüppell’s vultures (Gyps rueppellii)"

**File name:** SOM Social structure Rüppell’s Vultures.docx

**Supplementary Online Material**

**Table S1**. Overview of study subjects. …2
**Table S2**. Ethogram of agonistic interactions during food monopolization observations. …3
**Table S3**. Ethogram of social behaviour during focal observations. …3
**Table S4**. Overview of focal observation sampling effort. …5
**Appendix S1**. Inter-observer reliability test results. …6
**Table S5**. Hierarchy steepness based on Bayesian Elo-ratings. …7
**Appendix S2**. Formulas of models 1-5 and notes on model fit. …8
**Figure S1**. DHARMa residual diagnostic plots of model 1. …9
**Figure S2**. Model residuals plotted against each predictor variable of model 1. …10
**Figure S3**. DHARMa residual diagnostic plots of model 2. …11
**Figure S4**. Model residuals plotted against each predictor variable of model 2. …12
**Figure S5**. DHARMa residual diagnostic plots of model 3. …13
**Figure S6**. Model residuals plotted against each predictor variable of model 3. …14
**Figure S7**. DHARMa residual diagnostic plots of model 4. …15
**Figure S8**. Model residuals plotted against each predictor variable of model 4. …16
**Figure S9**. DHARMa residual diagnostic plots of model 5. …17
**Figure S10**. Model residuals plotted against each predictor variable of model 5. …18
**Table S6**. Summary table of outcome of conditional model 1. …19
**Table S7**. Summary table of outcome of conditional model 2. …19
**Table S8**. Summary table of outcome of conditional model 3. …19
**Table S9**. Summary table of outcome of conditional model 4. …19
**Table S10**. Summary table of outcome of conditional model 5. …20
**Table S11.** Summary table of significant outcomes of post-hoc tests of model 5. …20

Preprint - Not peer reviewed

***Table S1*** *Overview of study subjects. Individual characteristics of Rüppell's vultures included as subjects in this study (n = 23). Pair-bonds were already established before the start of the study, maintained a nest site throughout the study period, and incubated an (artificial) egg both breeding seasons. For offspring raised during the study period, the IDs of the pair that raised the offspring are provided. Periods in grey indicate that the individual was not included in analyses related to social bonds during that period. Age classes are based on birth year, determined at the start of each observation period. Immatures are individuals from their first non-breeding season onwards up to 4 years old (YO), young adults are between 5-10 YO, and adults are over 10 YO, (Bosè & Sarrazin 2007, Acácio* et al. *2024).*

| ID | Sex | Partner ID | (Foster) Parent IDs | Birth year | 2023 Breeding Jan-Jun | 2023 Non-breeding Jul-Dec | 2024 Breeding Jan-Jun | 2024 Non-breeding Jul-Dec |
|---|---|---|---|---|---|---|---|---|
| E | F | N | | ~95 | Adult | | | |
| L | M | M | | < 00 | Adult | | | |
| M | M | L | | < 00 | Adult | | | |
| N | M | E | | < 00 | Adult | | | |
| O | M | | | < 00 | Adult | | | |
| P | M | | | < 00 | Adult | | | |
| Q | M | G | | < 00 | Adult | | | |
| R | M | F | | < 00 | Adult | | | |
| S | M | | | < 00 | Adult | | | |
| T | M | J | | < 00 | Adult | | | |
| F | F | R | | 09 | Adult | | | |
| G | F | Q | | 12 | Adult | | | |
| J | F | T | | 18 | Young adult | | | |
| H | M | | | 19 | Immature | | Young adult | |
| I | F | | | 19 | Immature | | Young adult | |
| K | F | | | 22 | Immature | | Removed | |
| A | M | | L, M | 23 | Chick | | Immature | |
| B | M | | E, N | 23 | Chick | Immature | | Removed |
| C | F | | J, T | 23 | Chick | | Immature | |
| D | M | | G, Q | 23 | Chick | | Immature | |
| V | F | | F, R | 23 | Chick | | Immature | |
| U | F | | G, Q | 24 | | | Chick | Immature |
| W | M | | F, R | 24 | | | Chick | Immature |

Preprint - Not peer reviewed

***Table S2*** *Ethogram of agonistic interactions recorded during food monopolization observations.*

| Agonistic behaviour | Definition |
|---|---|
| Displacement | One bird enters in proximity with another bird who retreats / moves away within a few seconds and takes its position without any other agonistic behaviour. The actor of the displacement is considered to have won the interaction. |
| Chase | One bird pursues another one, typically flying. The actor of the chase is considered to have won the interaction. |
| Threat | One bird threatens another one by displaying an attack in its direction without touching it or by adopting a dominant posture toward the victim, most often by opening its wings with grunts or whistles. |
| Peck | One bird hits another one with its beak, it implies physical contact. |
| Kick | One bird hits another one with its foot, it implies physical contact. |
| Jump | One bird jumps on another's back and crushes it with all its weight, it implies physical contact. |
| Fight | Two birds hit each other intensely with their feet or beaks, with one or both birds jumping in the air. |
| Pull feathers | One bird pulls the feathers of another one with its beak. |
| Responses to agonistic behaviour | Definition |
| Retreat | One bird moves more than one meter away after receiving aggression from another bird. |
| Submissive response | One bird displaying a submissive posture (e.g., steps back, leans forward) or leaving its place ( < 1 meter) after receiving aggression from another bird. |

***Table S3*** *Ethogram of social behaviour recorded during focal observations.*

| Behaviour | Definition | Measure |
|---|---|---|
| Proximity | One birds' body is within one wing-length of another birds' body for at least 3 seconds. | Minutes per hour observed (m/h) |
| Staying on the nest | One bird is within the perimeter of a nest for at least 3 seconds. For nests on the ground, the perimeter is the area within one wing-length of the nest composed of straw and branches. For nests in nest-boxes, the perimeter includes the | Minutes per hour observed (m/h) |

Preprint - not peer reviewed

| | |
|---|---|
| | inside of the nest-box and the perches that are parallel to and directly attached to the nestbox. |

Preprint - Not peer reviewed

***Table S4*** *Overview of focal observation sampling effort. Sampling effort for focal observations of all subjects and all periods, excluding time out of sight and including or excluding time focal spend on a nest. Cells highlighted in grey indicate individual was absent during a period either because of young age or because of moving out of the group.*

| | Total observed time (hh:mm) | | | | Observed time off nest (hh:mm) | | | |
|---|---|---|---|---|---|---|---|---|
| | 2023 | | 2024 | | 2023 | | 2024 | |
| ID | Breeding | Non-breeding | Breeding | Non-breeding | Breeding | Non-breeding | Breeding | Non-breeding |
| E | 05:10 | 05:10 | 11:57 | 07:50 | 03:30 | 05:01 | 06:08 | 06:46 |
| L | 04:30 | 05:00 | 11:57 | 05:27 | 03:54 | 04:18 | 06:43 | 02:26 |
| M | 05:10 | 05:10 | 16:26 | 06:07 | 02:30 | 03:20 | 10:14 | 03:49 |
| N | 05:00 | 04:50 | 11:49 | 07:40 | 04:15 | 04:35 | 06:18 | 06:59 |
| O | 04:40 | 04:10 | 11:38 | 07:56 | 04:40 | 04:10 | 11:38 | 07:56 |
| P | 04:50 | 03:50 | 06:16 | 08:20 | 04:49 | 03:34 | 06:00 | 07:49 |
| Q | 05:10 | 05:00 | 11:49 | 10:25 | 03:16 | 03:59 | 04:35 | 08:09 |
| R | 04:50 | 04:50 | 11:50 | 05:21 | 02:53 | 03:31 | 04:50 | 02:26 |
| S | 04:50 | 04:50 | 11:53 | 05:40 | 04:50 | 04:50 | 11:53 | 05:40 |
| T | 05:00 | 05:00 | 11:50 | 09:21 | 04:30 | 04:11 | 06:27 | 07:03 |
| F | 05:00 | 05:00 | 12:04 | 05:16 | 02:54 | 02:42 | 03:43 | 02:45 |
| G | 05:20 | 05:10 | 12:05 | 05:44 | 03:30 | 02:51 | 04:08 | 03:10 |
| J | 05:10 | 05:00 | 13:23 | 07:00 | 02:21 | 03:39 | 09:29 | 05:02 |
| H | 05:00 | 04:40 | 12:18 | 06:40 | 04:56 | 04:26 | 11:54 | 06:21 |
| I | 04:50 | 05:10 | 12:14 | 04:29 | 04:41 | 05:10 | 12:08 | 04:29 |
| K | 05:10 | 05:10 | 11:14 | | 05:09 | 05:10 | 11:13 | |
| A | | 05:00 | 12:33 | 05:52 | | 04:18 | 12:29 | 05:52 |
| B | | 04:00 | 11:07 | | | 03:52 | 11:05 | |
| C | | 03:50 | 12:00 | 05:41 | | 03:50 | 12:00 | 05:39 |
| D | | 05:10 | 12:33 | 05:40 | | 04:40 | 12:13 | 05:40 |
| V | | 04:10 | 11:54 | 07:06 | | 03:00 | 11:46 | 07:06 |
| U | | | | 04:32 | | | | 03:38 |
| W | | | | 06:26 | | | | 06:26 |

Preprint not peer reviewed

**Appendix S1** *Inter-observer reliability test results.*

Coding of agonistic interactions during feedings was done by three different observers, E.S.J.v.D., S.W., and A.B., each coding a separate set of feeding videos. Coders achieved a good inter-rater reliability (ICC(A,1) = 0.784, 95%CI [0.746;0.817]) (Koo & Li 2016) for number of interactions per directional dyads per feeding observation (n = 337 across 3 feedings).

E.S.J.v.D., S.W., J.H.S., J.Z., Z.T., A.B., M.H.L.H. also each coded a different set of focal observations, scoring social proximity (within one wing-length) involving the focal subject and time on the nest. E.S.J.v.D., S.W., Z.T., A.B., M.H.L.H. had a good interrater reliability (ICC(A,1) = 0.792, 95%CI [0.73, 0.847]) (Koo & Li 2016) across all 62 focal observations. The work of J.H.S. and J.Z. could not be compared directly as they did not code the same subset of videos. E.S.J.v.D., S.W., Z.T., A.B., M.H.L.H. and J.Z. had a good inter rater reliability (ICC(A,1) = 0.821, 95%CI [0.741, 0.888]) (Koo & Li 2016) when comparing the duration of proximity and/or staying in the nest per focal observation (n = 40 where at least one coder observed proximity or staying in the nest, across 29 focal observations). E.S.J.v.D., S.W., Z.T., A.B., M.H.L.H. and J.H.S. had a moderate interrater reliability (ICC(A,1) = 0.707, 95%CI [0.603, 0.803]) (Koo & Li 2016) when comparing the duration of proximity and/or staying in the nest per focal observation (n = 45 across 33 focal observations).

Preprint - Not peer reviewed

***Table S5*** *Hierarchy steepness based on Bayesian Elo-ratings. Information on the food monopolization observations, recorded agonistic interactions with a clear winner and the steepness inferred from the Bayesian Elo-ratings calculated using these interactions.*

| | 2023 | | 2024 | |
|---|---|---|---|---|
| | Breeding | Non-breeding | Breeding | Non-breeding |
| N observations | 12 | 6 | 13 | 9 |
| First – last date (DD/MM) | 22/03 - 08/06 | 19/09 - 08/12 | 06/02 – 25/06 | 16/07 – 13/12 |
| N individuals | 16 | 21 | 21 | 21 |
| Decided agonistic interactions | 1304 | 1265 | 5770 | 876 |
| Interactions per individual | 81.5 | 60.2 | 274.8 | 41.7 |
| Interactions per dyad | 10.9 | 6.0 | 27.5 | 4.2 |
| Proportion unknown dyads | 0.025 | 0.219 | 0.024 | 0.229 |
| median k value estimate (89% CI) | 0.16 (0.10-0.22) | 0.25 (0.18-0.33) | 0.25 (0.21-0.30) | 0.19 (0.09-0.29) |
| Mean steepness (89% CI) | 0.58 (0.54-0.61) | 0.59 (0.57-0.60) | 0.74 (0.74-0.75) | 0.50 (0.45-0.55) |

Preprint - Not peer reviewed

**Appendix S2.** *Formulas of models 1-5 and notes on model fit for dominance rank and social bond analyses.*

**Model 1:** GLMM with gaussian (log link ) family of z-transformed Elo score.

Elo score ~ Sex + Age class + Pair-bond status * Season + (1 | individual id) + (1 | Period)

Model diagnostics of model 1 (Figure S1, Figure S2) suggested a good fit.

**Model 2:** GLMM with gamma (log link) family of strength.

Strength ~ Pair-bond status * Season + Sex + Age + Mean Elo-score z-transformed + (1 | Period) + (1 | Name)

Model diagnostics of model 2 (Figure S3, Figure S4) suggested a good fit.

**Model 3:** GLMM with gamma (log link) family of Eigenvector centrality.

Eigenvector centrality ~ Pair-bond status * Season + Sex + Age + Mean Elo-score z-transformed + (1 | Period) + (1 | Name)

Model diagnostics of model 3 (Figure S5, Figure S6) suggested a generally good fit but did indicate minor deviations in the upper quantile. Residuals showed no patterns against most predictors, but the z-transformed mean Elo score did show deviations in the upper quantile. As the combined adjusted quantile test for this was not significant and the deviation itself was minimal, we concluded that the model was correctly specified (Hartig *et al.* 2024).

**Model 4:** GLMM with gamma (log link) family of number of strong bonds.

Strong_ties ~ Pair-bond status * Season + Sex + Age + (1 | Period) + (1 | Name)

Model diagnostics of model 4 (Figure S7, Figure S8) suggested a good fit as the residuals met model assumptions and no quantile deviations were detected for residuals plotted against predicted values or against each predictor separately.

**Model 5:** GLMM with Tweedie (log link) family of proximity (m/h) per dyad.

Proximity (m/h) ~ Dyadic sex combination + Dyadic age combination + Difference in Elo-score, z-transformed + Season * Pair-bond status + (1 | Period) + (1 | Dyad_id)

Model diagnostics of model 5 (Figure S9, Figure S10) suggested a generally good fit and residuals showed no patterns against most predictors but did indicate homogeneity of variance between the two seasons and between dyadic age class combinations.

Preprint - Not Peer-reviewed

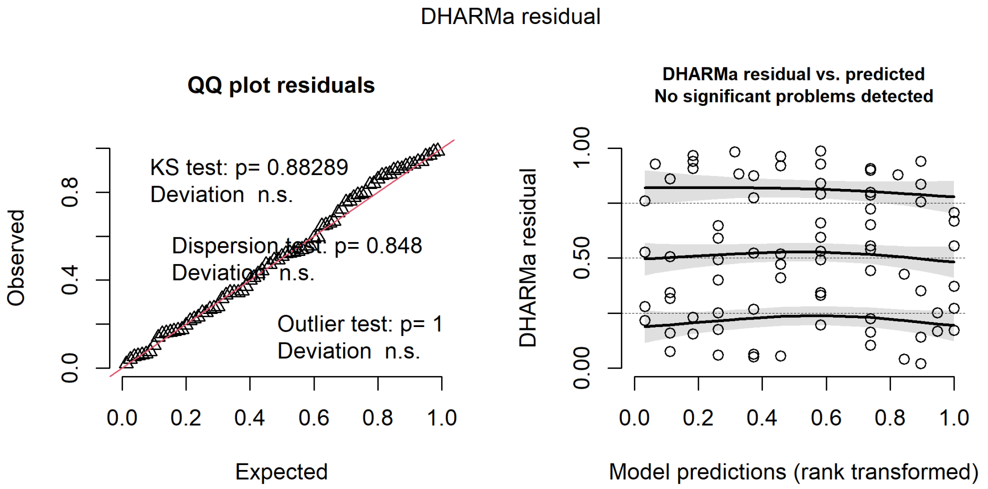

***Figure S1.*** *DHARMa residual diagnostic plots of model 1. QQ-plot with added tests for correct distribution, dispersion, and outliers (left). Residuals plotted against the predicted variable (right) with .25, .5 and .75 quantile regression lines.*

Preprint - Not peer reviewed

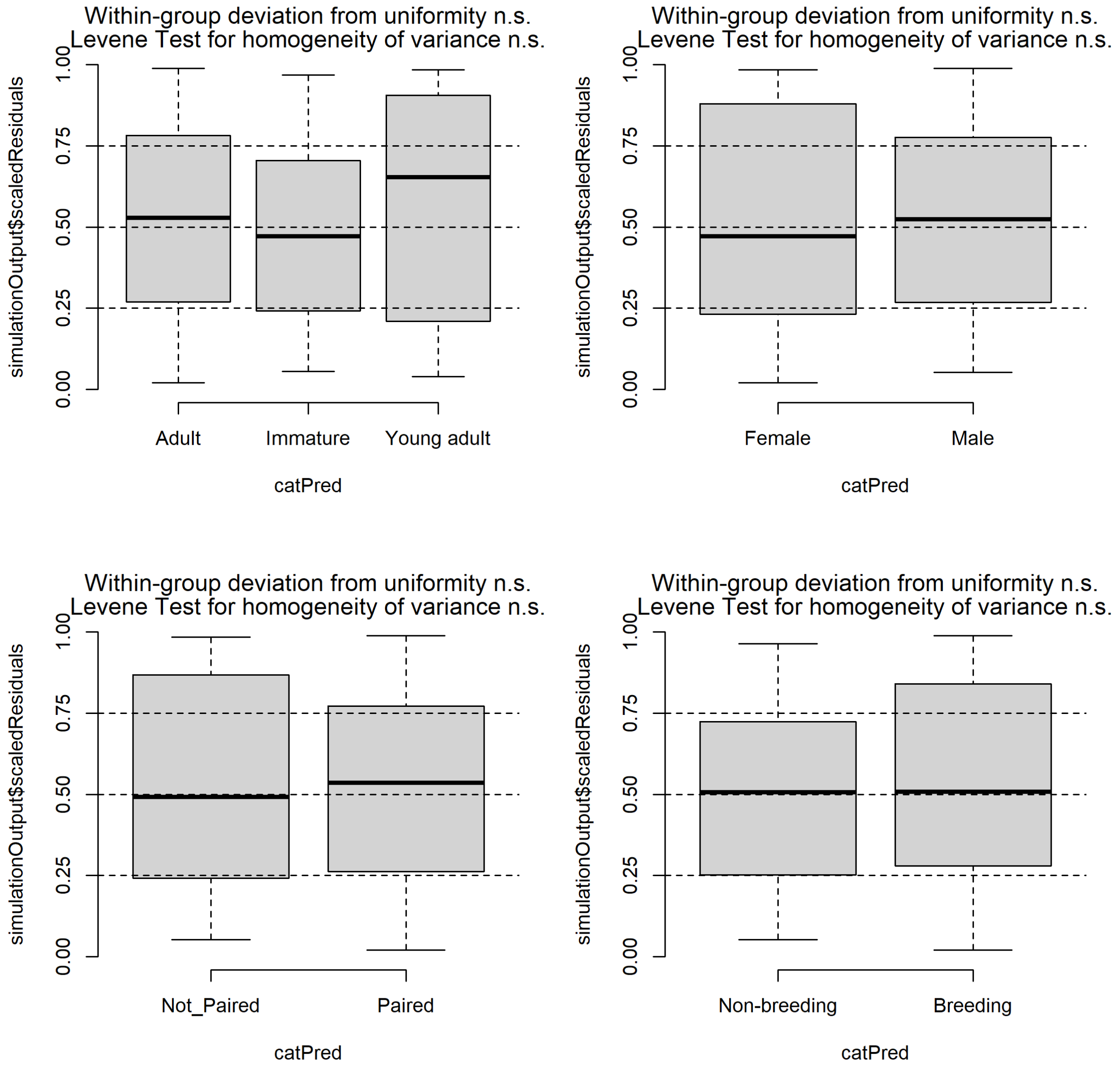

***Figure S2.*** *Model residuals plotted against each separate predictor variable of model 1. Age class (top left), sex (top right), pair-bond status (bottom left), season (bottom right) with the outcome of a test for uniformity per level of the categorical predictors and a test of homogeneity of variance between levels per categorical predictor.*

Preprint

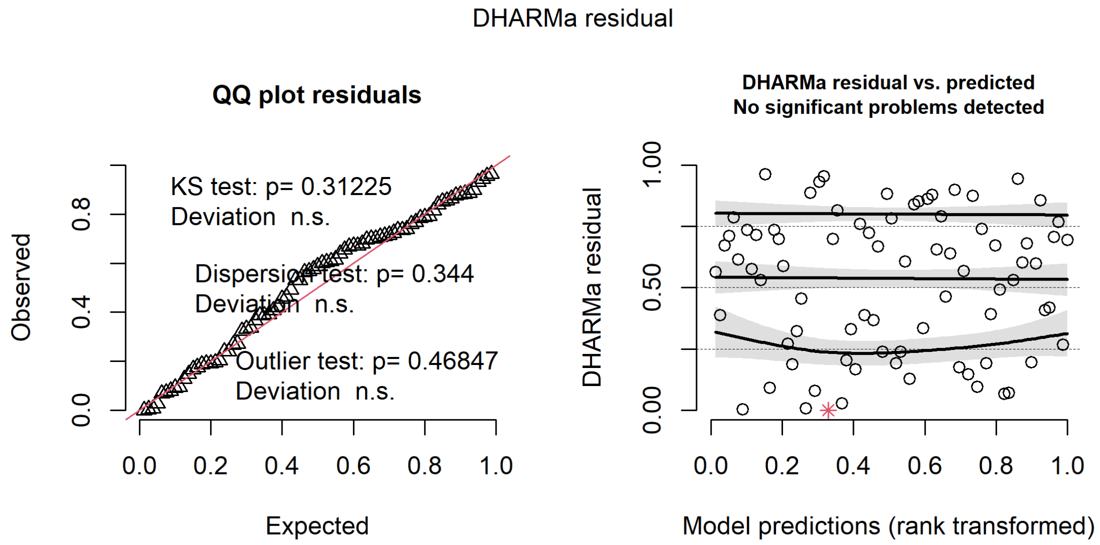

***Figure S3**. DHARMa residual diagnostic plots of model 2. QQ-plot with added tests for correct distribution, dispersion, and outliers (left). Residuals plotted against the predicted variable (right) with .25, .5 and .75 quantile regression lines.*

Preprint - Not peer rev

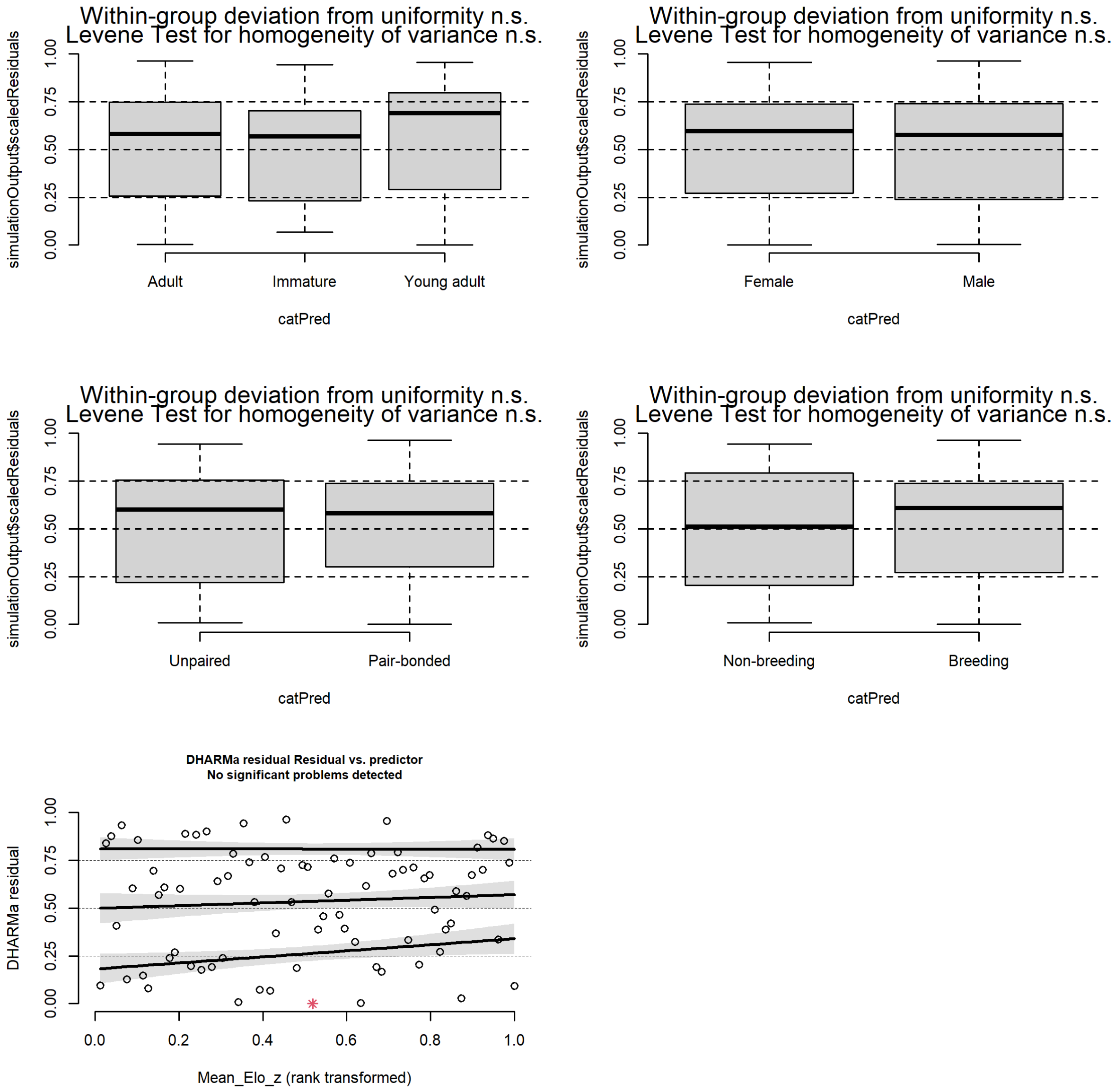

***Figure S4.*** *Model residuals plotted against each separate predictor variable of model 2. Age class (top left), sex (top right), pair-bond status (middle left), season (middle right), mean z-transformed Elo-score (bottom left) with the outcome of a test for uniformity per level of the categorical predictors and a test of homogeneity of variance between levels per categorical predictor.*

Preprint

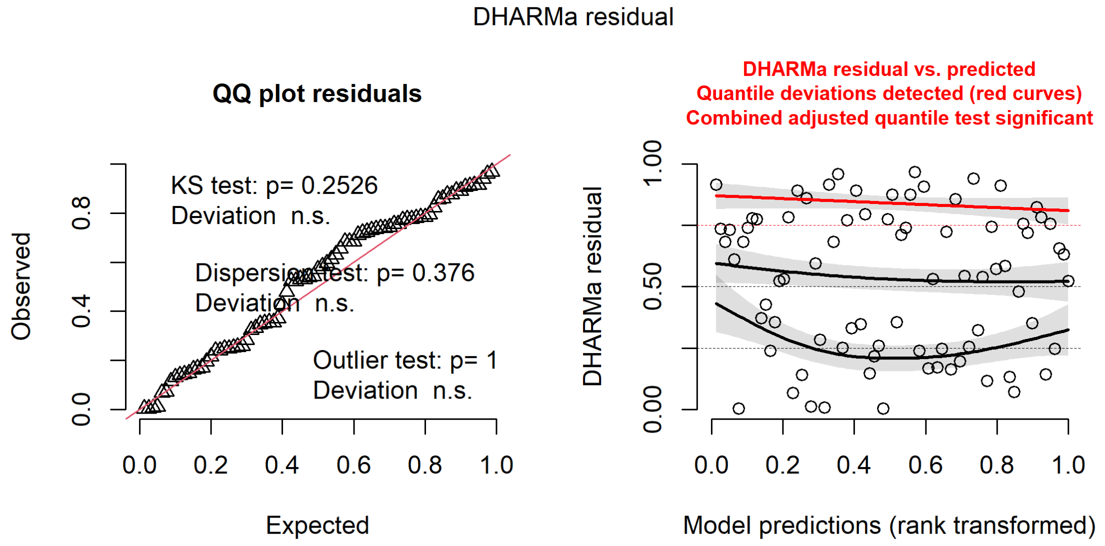

***Figure S5.*** *DHARMa residual diagnostic plots of model 3. QQ-plot with added tests for correct distribution, dispersion, and outliers (left). Residuals plotted against the predicted variable (right) with .25, .5 and .75 quantile regression lines.*

Preprint - Not peer r

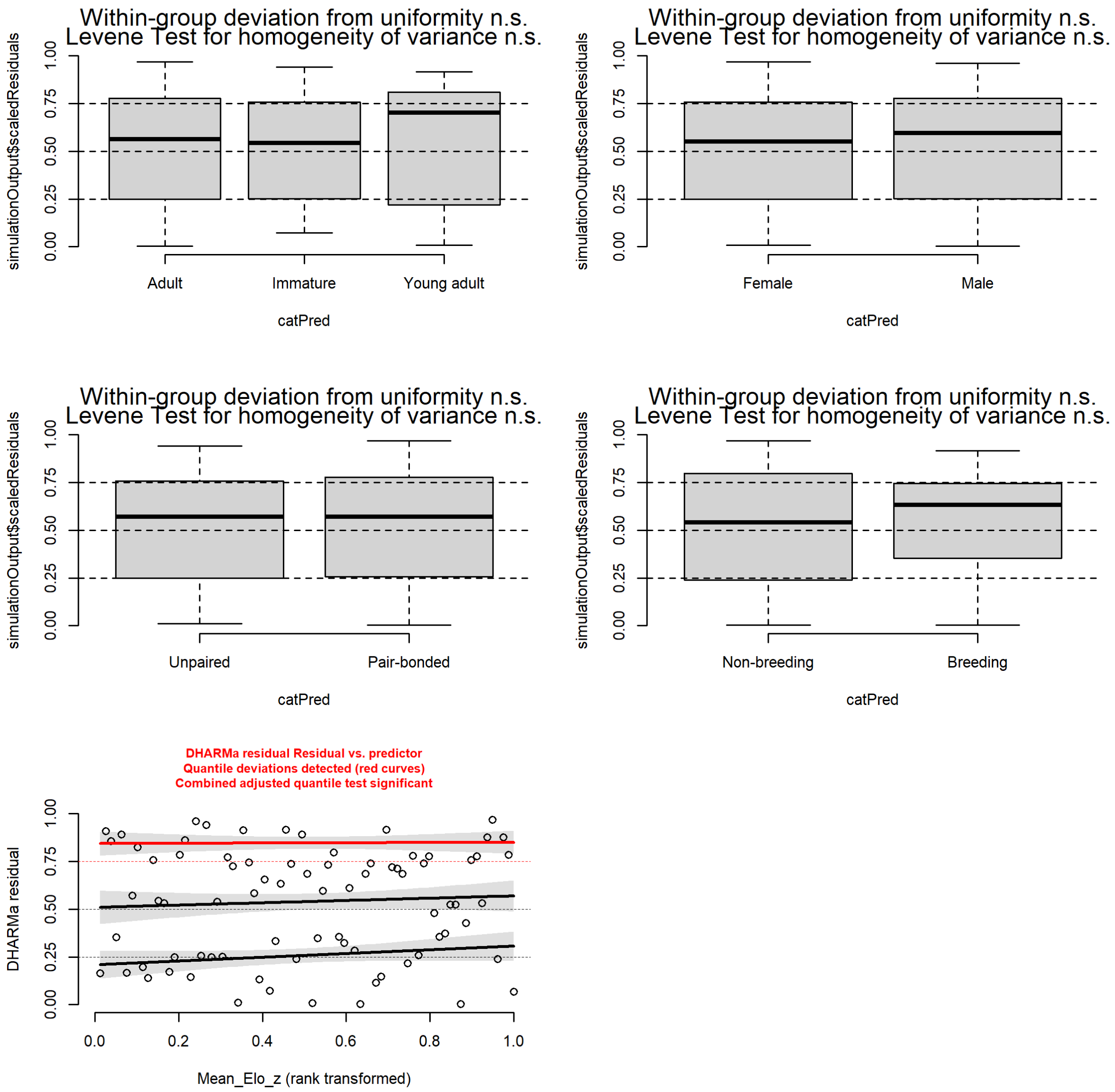

***Figure S6.*** *Model residuals plotted against each separate predictor variable of model 3. Age class (top left), sex (top right), pair-bond status (middle left), season (middle right), mean z-transformed Elo-score (bottom left) with the outcome of a test for uniformity per level of the categorical predictors and a test of homogeneity of variance between levels per categorical predictor.*

Preprint

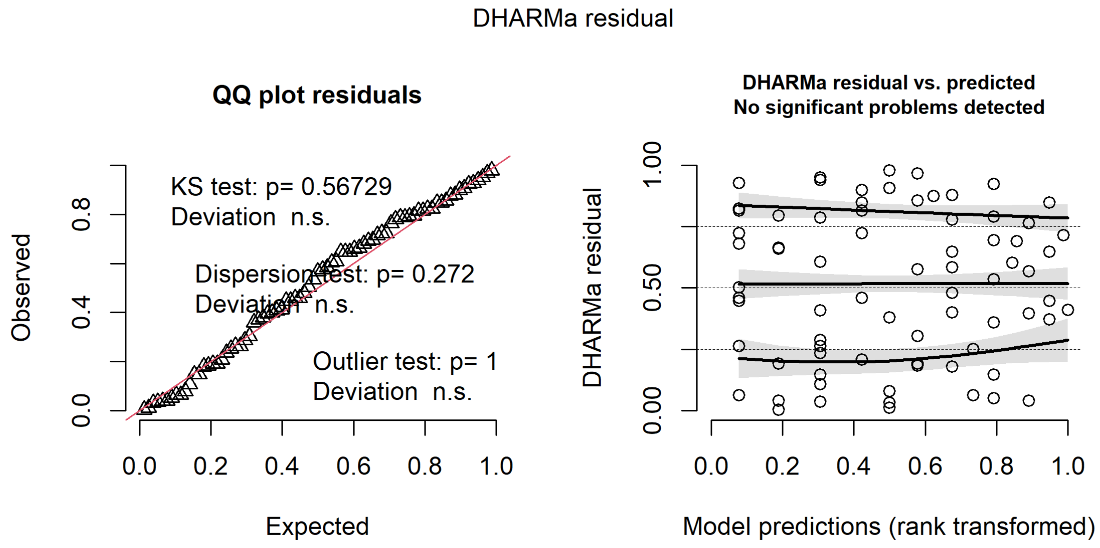

***Figure S7***. *DHARMa residual diagnostic plots of model 4. QQ-plot with added tests for correct distribution, dispersion, and outliers (left). Residuals plotted against the predicted variable (right) with .25, .5 and .75 quantile regression lines.*

Preprint - Not peer reviewed

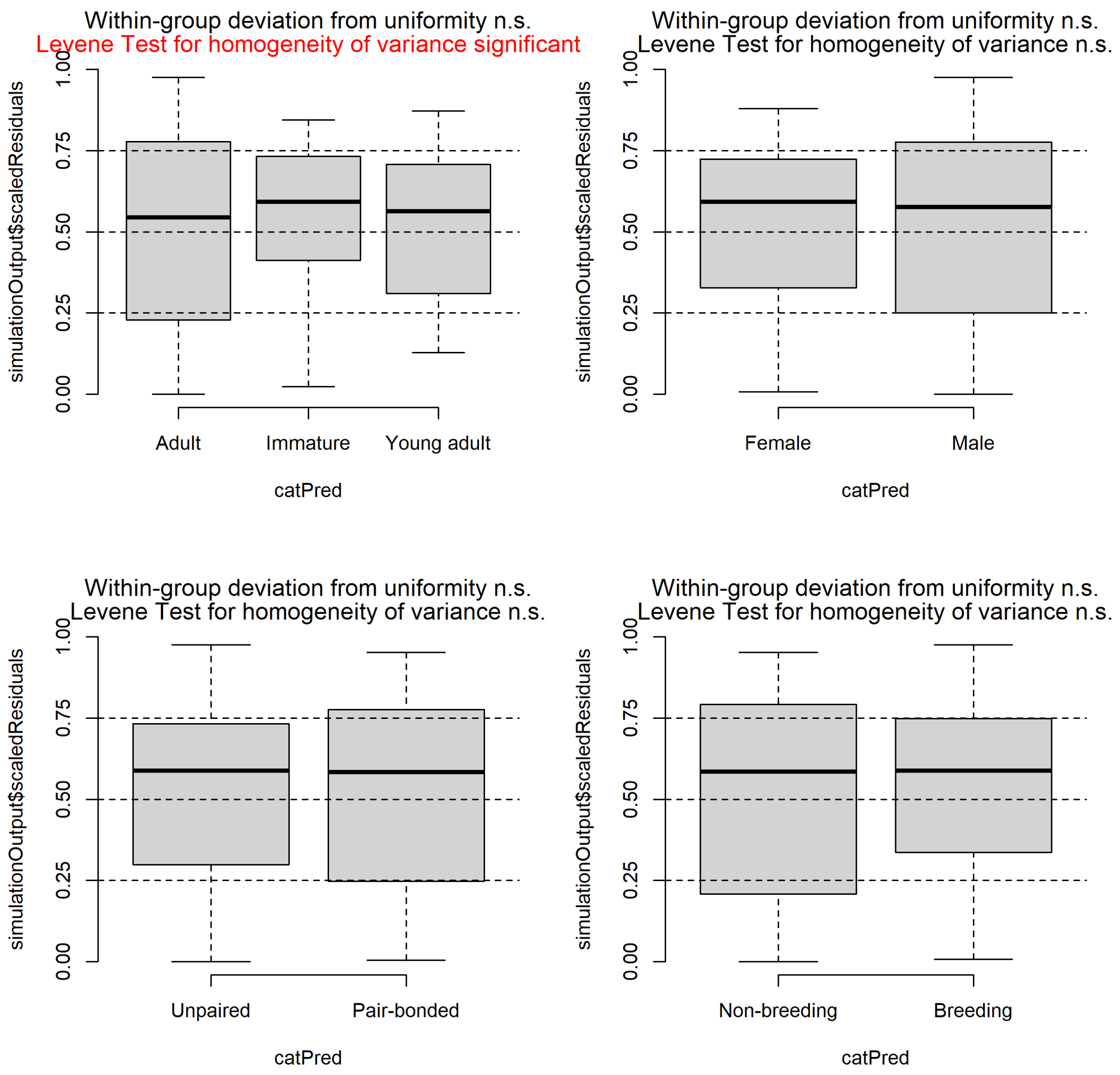

***Figure S8.*** *Model residuals plotted against each separate predictor variable of model 4. Age class (top left), sex (top right), pair-bond status (bottom left), season (bottom right) with the outcome of a test for uniformity per level of the categorical predictors and a test of homogeneity of variance between levels per categorical predictor.*

Preprint

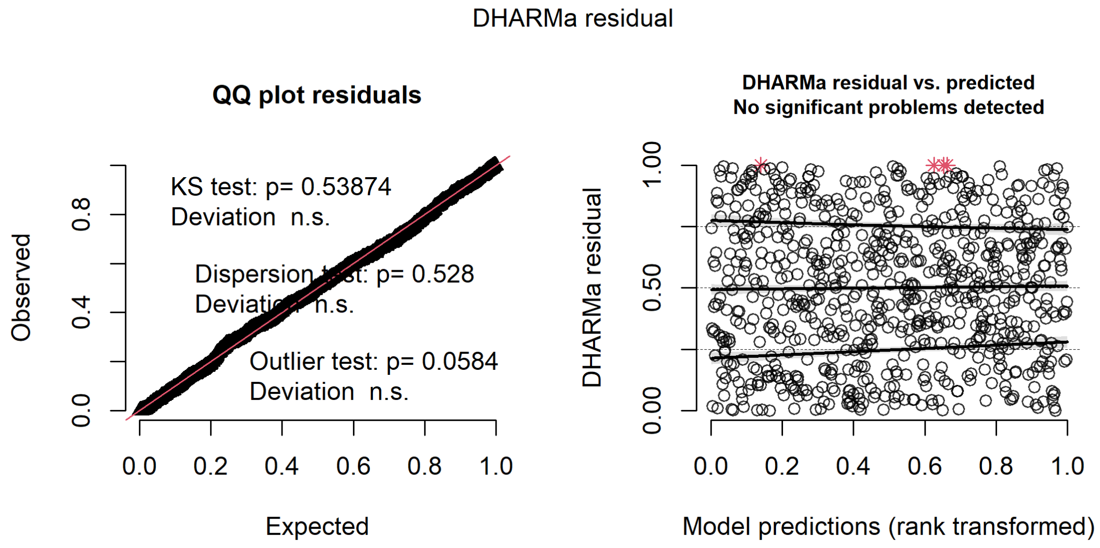

***Figure S9.*** *DHARMa residual diagnostic plots of model 5. QQ-plot with added tests for correct distribution, dispersion, and outliers (left). Residuals plotted against the predicted variable (right) with .25, .5 and .75 quantile regression lines.*

Preprint - Not peer reviewed

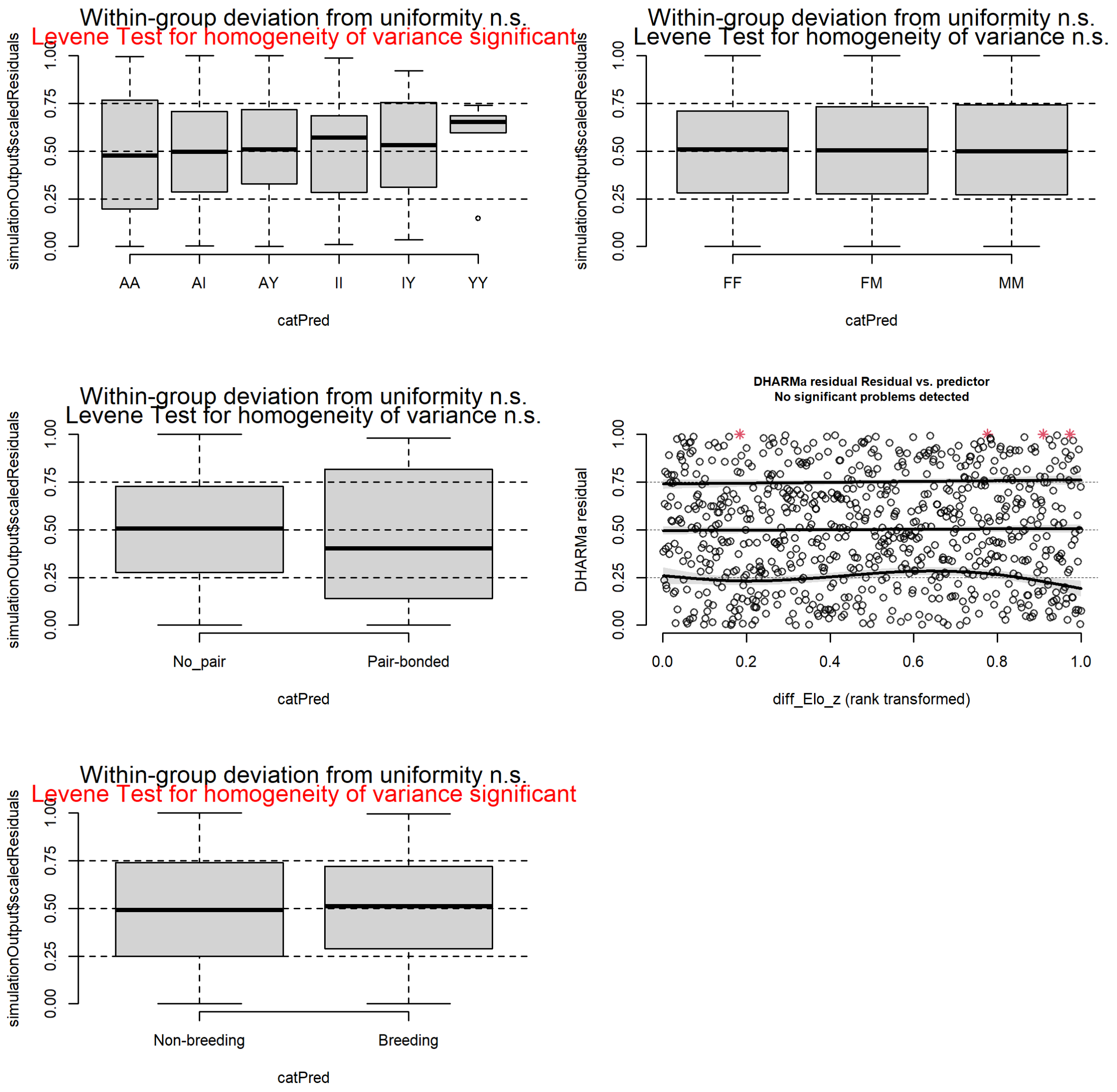

***Figure S10.*** *Model residuals plotted against each separate predictor variable of model 5. Dyadic age combination (top left) with labels corresponding to the combination of A=adult, Y=young adult and I=immature. Dyadic sex (top right) with FF=female-female, MM=male-male and FM=female-male, pair-bond status (middle left), z-transformed difference in Elo-score (middle right) and season (bottom left). Each plot with the outcome of a test for uniformity per level of the categorical predictors and a test of homogeneity of variance between levels per categorical predictor.*

Preprint

***Table S6.*** *Summary table of outcome of conditional model 1. GLMM with gaussian (log link) family of z-transformed Elo-score. Significance codes: *** 0.001, ** 0.01, * 0.05, . 0.1.*

| | Estimate | Std. Error | z value | Pr( > \|z\|) | |
|---|---|---|---|---|---|
| (Intercept) | 2.474 | 0.167 | 14.856 | < 0.001 | *** |
| Male | -0.33 | 0.086 | -3.822 | < 0.001 | *** |
| Immature | -0.25 | 0.129 | -1.937 | 0.053 | . |
| Young adult | -0.004 | 0.13 | -0.027 | 0.978 | |
| Pair-bonded | 0.164 | 0.119 | 1.386 | 0.166 | |
| Breeding season | -0.441 | 0.165 | -2.67 | 0.008 | ** |
| Pair-bonded: Breeding season | 0.401 | 0.103 | 3.909 | < 0.001 | *** |

***Table S7.*** *Summary table of outcome of conditional model 2. GLMM with gamma (log link) family of strength. Significance codes: *** 0.001, ** 0.01, * 0.05, . 0.1.*

| | Estimate | Std. Error | z value | Pr( > \|z\|) | |
|---|---|---|---|---|---|
| (Intercept) | 3.677 | 0.141 | 26.039 | < 0.001 | *** |
| Pair-bonded | 0.128 | 0.124 | 1.028 | 0.304 | |
| Breeding season | 0.171 | 0.127 | 1.352 | 0.176 | |
| Male | -0.084 | 0.096 | -0.867 | 0.386 | |
| Immature | 0.251 | 0.119 | 2.105 | 0.035 | * |
| Young adult | 0.108 | 0.134 | 0.81 | 0.418 | |
| Mean Elo-score z-transformed | -0.001 | 0.056 | -0.021 | 0.983 | |
| Pair-bonded: Breeding season | -0.48 | 0.139 | -3.442 | 0.001 | *** |

***Table S8.*** *Summary table of outcome of conditional model 3. GLMM with gamma (log link) family of Eigenvector centrality. Significance codes: *** 0.001, ** 0.01, * 0.05, . 0.1.*

| | Estimate | Std. Error | z value | Pr( > \|z\|) | |
|---|---|---|---|---|---|
| (Intercept) | -0.621 | 0.142 | -4.37 | < 0.001 | *** |
| Pair-bonded | 0.095 | 0.134 | 0.708 | 0.479 | |
| Breeding season | 0.214 | 0.103 | 2.074 | 0.038 | * |
| Male | -0.086 | 0.105 | -0.819 | 0.413 | |
| Immature | 0.266 | 0.13 | 2.041 | 0.041 | * |
| Young adult | 0.077 | 0.143 | 0.535 | 0.593 | |
| Mean Elo-score z-transformed | 0.008 | 0.06 | 0.132 | 0.895 | |
| Pair-bonded: Breeding season | -0.461 | 0.145 | -3.172 | 0.002 | ** |

***Table S9.*** *Summary table of outcome of conditional model 4. GLMM with gamma (log link) family of number of strong bonds. Significance codes: *** 0.001, ** 0.01, * 0.05, . 0.1.*

| | Estimate | Std. Error | z value | Pr( > \|z\|) | |
|---|---|---|---|---|---|
| (Intercept) | 1.777 | 0.182 | 9.745 | < 0.001 | *** |
| Pair-bonded | -0.047 | 0.172 | -0.274 | 0.784 | |
| Breeding season | 0.138 | 0.134 | 1.029 | 0.304 | |
| Male | -0.025 | 0.114 | -0.219 | 0.827 | |
| Immature | 0.376 | 0.163 | 2.305 | 0.021 | * |
| Young adult | 0.411 | 0.192 | 2.141 | 0.032 | * |
| Pair-bonded: Breeding season | -0.356 | 0.189 | -1.877 | 0.06 | . |

Preprint - Not peer reviewed

***Table S10.*** *Summary table of outcome of conditional model 5. GLMM with Tweedie (log link) family of dyadic proximity rate (m/h). Significance codes: *** 0.001, ** 0.01, * 0.05, . 0.1.*

| | Estimate | Std. Error | z value | Pr( > \|z\|) | |
|---|---|---|---|---|---|
| (Intercept) | 0.417 | 0.135 | 3.088 | 0.002 | ** |
| Female-Male | -0.042 | 0.096 | -0.441 | 0.659 | |
| Male-Male | -0.103 | 0.105 | -0.984 | 0.325 | |
| Adult-Immature | 0.243 | 0.08 | 3.053 | 0.002 | ** |
| Adult-Young adult | 0.042 | 0.11 | 0.384 | 0.701 | |
| Immature-Immature | 0.944 | 0.113 | 8.369 | < 0.001 | *** |
| Immature-Young adult | 0.641 | 0.13 | 4.927 | < 0.001 | *** |
| Young adult - Young adult | 0.291 | 0.325 | 0.895 | 0.371 | |
| Difference Elo-score z-transformed | 0.045 | 0.03 | 1.505 | 0.132 | |
| Breeding season | 0.197 | 0.121 | 1.625 | 0.104 | |
| Pair-bonded | 2.061 | 0.193 | 10.686 | < 0.001 | *** |
| Breeding season: Pair-bonded | -1.191 | 0.248 | -4.81 | < 0.001 | *** |

***Table S11 .*** *Summary table of significant (p < 0.05) outcomes of post-hoc tests comparing different age class combinations of model 5. GLMM with Tweedie (log link) family of dyadic proximity rate (m/h). Significance codes: *** 0.001, ** 0.01, * 0.05, . 0.1.*

| Dyadic age class combination | Estimate | Std. Error | z value | Pr( > \|z\|) | |
|---|---|---|---|---|---|
| AA - AI | -0,243 | 0,08 | -3,053 | 0,027 | * |
| AA - II | -0,944 | 0,113 | -8,369 | < 0.001 | *** |
| AA - IY | -0,641 | 0,13 | -4,927 | < 0.001 | *** |
| AI - II | -0,701 | 0,105 | -6,686 | < 0.001 | *** |
| AI - IY | -0,398 | 0,122 | -3,251 | 0,015 | * |
| AY - II | -0,902 | 0,128 | -7,033 | < 0.001 | *** |

Preprint - Not peer reviewed
